# Protein citrullination was introduced into animals by horizontal gene transfer from cyanobacteria

**DOI:** 10.1101/2020.06.13.150037

**Authors:** Thomas F. M. Cummings, Kevin Gori, Luis Sanchez-Pulido, Gabriel Gavriilidis, David Moi, Abigail R. Wilson, Elizabeth Murchison, Christophe Dessimoz, Chris P. Ponting, Maria A. Christophorou

## Abstract

Protein post-translational modifications (PTMs) add an enormous amount of sophistication to biological systems but their origins are largely unexplored. Citrullination, a key regulatory mechanism in human physiology and pathophysiology, is particularly enigmatic in an evolutionary context. The citrullinating enzymes peptidylarginine deiminases (PADIs) are ubiquitous across vertebrates but absent from yeast, worms and flies. Here, we map the surprising evolutionary trajectory of PADIs into the animal lineage. We present strong phylogenetic support for a clade encompassing animal and cyanobacterial *PADIs* that excludes fungal and other bacterial homologues. The animal and cyanobacterial PADIs share unique, functionally relevant synapomorphies that are absent from all other homologues. Molecular clock calculations and sequence divergence analyses using the fossil record estimate the last common ancestor of the cyanobacterial and animal PADIs to be approximately 1 billion years old, far younger than the 3.35-4.52 billion years known to separate bacterial and eukaryotic lineages. Under an assumption of vertical descent, PADI sequence change is anachronistically slow during this evolutionary time frame, even when compared to mitochondrial proteins, products of likely endosymbiont gene transfer and some of the most highly conserved proteins in life. The consilience of evidence indicates that *PADIs* were introduced from cyanobacteria into animals by horizontal gene transfer (HGT). The ancestral cyanobacterial protein is enzymatically active and can citrullinate eukaryotic proteins, suggesting that the *PADI* HGT event introduced a new catalytic capability into the regulatory repertoire of animals. This study reveals the unusual evolution of a pleiotropic protein modification with clear relevance in human physiology and disease.

## Introduction

Post-translational modifications (PTMs) allow for temporal and spatial control of protein function in response to cellular and environmental signals and comprise an integral part of cellular and organismal life. The development of ever more sensitive and quantitative analytical methods has made possible the identification of PTMs within cells and has enhanced our understanding of the molecular and cellular functions they regulate. This has led to renewed interest in studying previously known, as well as newly identified modifications. Although PTMs have been classically studied in eukaryotic organisms, an increasing number of them are also discovered in bacteria^1,2^. However, very few of the >200 currently known PTMs have been studied in an evolutionary context. Some PTMs, such as phosphorylation, acetylation and glycosylation are ubiquitous across all domains of life suggesting that the enzymes that catalyse them existed in the Last Universal Common Ancestor (LUCA)^3^. In other cases, such as protein ubiquitylation, evolutionary analyses show that the catalyzing enzymes only appear in the last eukaryotic common ancestor, although domains analogous to E1 enzymes can also be found in bacteria^1,4^. Understanding the processes and adaptive constrains that have shaped the evolution of protein modifying enzymes can reveal which aspects are likely to be functionally important and may inform our understanding of their organismal roles.

Citrullination is the post-translational conversion of a protein arginine residue to the non-coded amino acid citrulline and is catalysed by PADIs enzymes in a calcium-dependent manner. Although citrullination involves a small mass change of only 0.98Da, the removal of a positive charge from the arginine side chain can lead to profound biochemical changes and is known to alter protein structure, sub-cellular localisation and affinity to other proteins and nucleic acids^5–11^. Via these alterations PADIs regulate fundamental physiological and cellular processes. The best-established role of citrullination is in innate immunity, through mediating the release of neutrophil extracellular traps (NETs)^12^ but a plethora of studies have shown that PADIs also regulate gene expression, chromatin compaction, nerve myelination, skin homeostasis and the establishment of ground state pluripotency^8,13,14^. Notably, deregulation of PADIs is strongly implicated in the aetiology of a host of pathologies including autoimmunity (rheumatoid arthritis, ulcerative colitis, psoriasis and type I diabetes), neurodegeneration (multiple sclerosis, Alzheimer’s and prion diseases) and metastatic cancer^13,15–18^. In the case of rheumatoid arthritis, autoantibodies against citrullinated endogenous proteins (Anti-Citrullinated Protein Antibodies, ACPAs) serve as diagnostic and prognostic markers as they precede the onset of symptoms by several years and correlate with disease severity and response to treatment^19^. Experimental *in vivo* models of PADI over-expression show that deregulation of citrullination drives the development of multiple sclerosis and cancer^16,20^, while genetic ablation or chemical inhibition of different PADI family members has been shown to mitigate against some the pathologies mentioned above^16,21–23^. Although the underlying mechanism in the above cases is aberrantly high citrullination, loss of PADI activity is also deleterious and has been shown to compromise neurodevelopment, fertility and embryo development^8,14,24^. PADIs have therefore emerged as important therapeutic targets^25^ and this has motivated the study of their exquisite regulation by calcium and the physiologically relevant mechanisms of their activation^26–28^.

In an evolutionary context, *PADIs* are puzzling. Orthologues of the human *PADIs* are ubiquitous in bony fish, birds, reptiles, amphibians and mammals, but are unexpectedly missing from many eukaryotes including plants, yeast, worms and insects. Indeed, their absence from these genetic model organisms has arguably impeded progress towards the understanding of their functions and citrullination has remained a rather obscure PTM for a long time. The *PADI* gene is widely thought to have appeared first in the last common ancestor of teleosteans and mammals^13,29,30^, with duplications in subsequent lineages resulting in five mammalian paralogues. Therefore, citrullination seemingly defies the perception that PTMs are of ancient origin.

Studying the evolutionary origin of PTM-catalyzing enzymes using protein sequence is particularly challenging as sequence similarity becomes less reliable with evolutionary distance, however structural homology can point to the evolutionary origins of proteins^3,31^. We therefore considered homology of PADI structural domains. Mammalian PADIs consist of three structural domains, the N-terminal (PAD_N, Pfam annotation: PF08526), middle (PAD_M, Pfam annotation: PF08527) and catalytic C-terminal domains (PAD_C, Pfam annotation: PF03068). Mammalian genomes encode two distant homologues of the PAD_C domain: N(G),N(G)-dimethylarginine dimethylaminohydrolase [DDAH] and Glycine amidinotransferase [AGAT]. Both, however, are divergent in sequence and lack PAD_N and PAD_M domains. Two other citrullinating enzymes are known among some bacteria and early-branching eukaryotes: pPAD, an extended agmatine deiminase found in *Porphyromonas gingivalis* and giardiaADI, an extended form of the free *L*-arginine deiminase gADI, found in the human parasite *Giardia Lamblia*^32,33^. These enzymes also lack PAD_N and PAD_M domains, are highly divergent in sequence and have different substrate specificities.

Although PADI proteins are widely considered to be specific to vertebrates, their crystal structures^28,34^ hint at a possibly more ancient origin as they reveal that the catalytic (PAD_C) domain adopts the same pentein fold as a variety of other widely distributed proteins that otherwise show little similarity in terms of amino acid conservation^35,36^ (Figure S1). These proteins include a broad family of guanidino-group (the functional group of the side chain of arginine and agmatine) modifying enzymes that possess hydrolase, dihydrolase and amidinotransferase catalytic activity, such as bacterial and early eukaryotic agmatine deiminases. The pentein-fold containing enzymes share a broad catalytic core of a Cys, His and two polar guanidine binding residues – Asp or Glu^36^. The presence of this ancient fold and catalytic triad within PAD_C suggests that it was present early in cellular life.

A 2015 study by Crisp *et al*., identified possible *PADI* homologues in some bacterial species. Based on the finding that a possible homologue could be identified in prokaryotes but not in multiple *Drosophila* and *Caenorhabditis* species, the authors included *PADIs* among a list of 145 genes proposed to have been transferred into the genome of a vertebrate ancestor of extant mammals by horizontal gene transfer (HGT, also known as lateral gene transfer)^37^. HGT is the non-heritable transmission of genetic material from one organism to another, often via a virus or mobile genetic element and involving endosymbiotic or commensal relationships between donor and recipient^38,39^. HGT is widespread among prokaryotes and is recognised as a mechanism that shapes the evolution and adaptive potential of bacteria, for example in the acquisition of antibiotic resistance^40,41^. Although many cases of horizontal transfer have been reported between bacteria and unicellular eukaryotes, fewer bacteria-to-animal HGT events have been studied to date ^38,42,43^. The majority of cases involve transfer into an invertebrate host, such as an insect or worm^44–48^, while HGT into animals with specialised germline cells is thought to be very rare^49^. Furthermore, these few accounts of bacteria-to-animal HGT have been the topic of intense debate^37,50–54^. Claims of HGT should therefore be considered on a case-by-case basis and tested against the alternative hypothesis of widespread gene loss^51^. In light of the absence of PADI homologues in most invertebrate animals, PADI evolution requires detailed consideration.

## Results

### Comprehensive Identification of PADI homologues

In order to understand the distribution and evolution of citrullination we sought to identify all *PADI* homologues from across life. We started by collecting orthologous *PADIs* using the EggNOG database, employing an unsupervised clustering algorithm of all proteins contained in 2031 genomes across cellular life^55^. To expand on this list, we used HMMER searches to identify all sequences in current sequence databases that contain a PAD_C domain, as defined by having significant sequence similarity (E-value < 1×10^−3^), and assessed these for the presence of critical substrate-binding and calcium-binding residues annotated to human PADIs^28^. This was supplemented by additional iterative jackhmmer searches and Position-Specific Iterated BLAST (PSI-BLAST) searches as well as tblastn searches of genomic databases.

The taxonomic distribution of *PADIs* and proportion of species that harbour a PADI orthologue are presented in Table 1. PADIs are not ubiquitous across the metazoa, but are found in all major branches of the vertebrates (jawless fish, sharks and rays, bony fish, amphibians, reptiles, birds and mammals). Out of all species whose genomes have been sequenced to date, the earliest diverging invertebrate animals with a *PADI* gene are *Priapulus caudatus* (an ecdysozoan), *Saccoglossus kowalevskii* (a hemichordate), and *Branchiostoma belcheri* (a cephalochordate). Surprisingly, we identified a large number of PADI sequences with conservation of substrate and calcium-binding residues in bacteria and fungi. *PADIs* are also not ubiquitous across bacteria, and are most prevalent within cyanobacteria. None of the known plants or animals diverging before opisthokonts have a detectable *PADI* homologue. Our searches also returned two outliers, one in archaea and one in viruses. However, upon closer inspection, both hits were determined to be due to misattribution (Figures S2, S3; *see also* Methods) and were therefore not included in further analyses. This taxonomic distribution could suggest an evolutionary model in which *PADI* genes were lost independently in many separate lineages. In this scenario, gene loss occurred in all early-branching lineages leading to at least 306 non-opisthokont eukaryotes and in other lineages, for example those leading to *Drosophila* and *Caenorhabditis*.

**Table 1:**
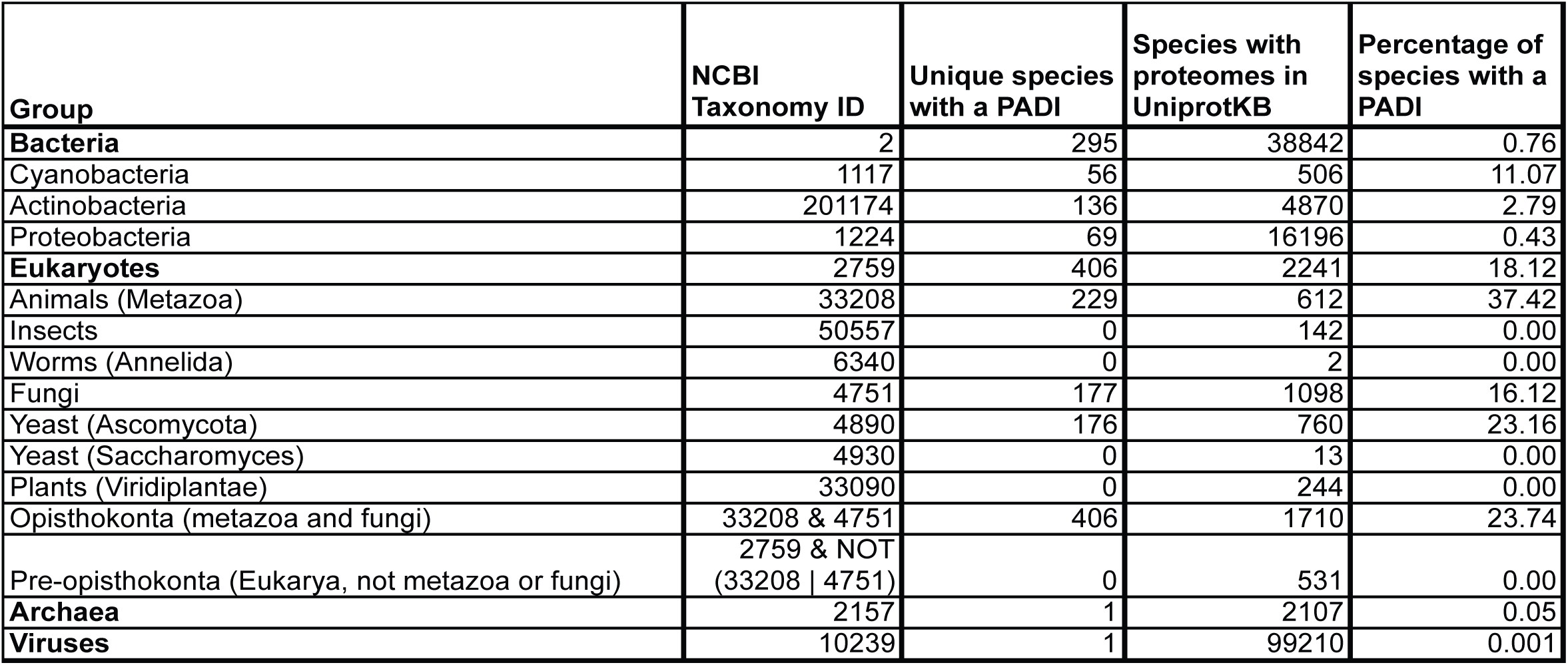
The number and proportion of species harbouring a putative PADI orthologue. HMM searches (https://www.ebi.ac.uk/Tools/hmmer) for similarity to the vertebrate PAD_C domain from human PADI2, were carried out using HmmerWeb version 2.41.1 against the UniProtKB (v.2019_09) database. Unique species with significant sequence similarity (E-value < 1×10-3) are presented. Proportions are given relative to the total number of species in within UniProtKB, for each group.

To test whether bacterial and fungal PADIs are related to pPAD and gADI enzymes, we conducted an unrooted maximum likelihood phylogenetic analysis using putative PADI homologues and sequences with significant HMMER similarity to pPAD and gADI sequences (Figure S4). Bacterial, fungal and animal PADIs form a single monophyletic outgroup that excludes both pPAD and gADI enzyme types, showing that each of the three types of protein is phyletically distinct. The pPAD and gADI type proteins can therefore be excluded from further consideration of the evolutionary origin of animal PADIs.

### A strongly supported phylogenetic clade contains cyanobacterial and animal but not fungal PADIs

We next performed detailed phylogenetic analyses. All bacterial PADI sequences within the Pathosystems Resource Integration Center (PATRIC) database were aligned with a subsampled set of PADI sequences from animals and fungi selected to maximise both protein sequence diversity and lineage representation. A maximum likelihood approach with the best fitting evolutionary rate model was used to produce an initial phylogenetic tree. Very strong bootstrap support (>95%) was obtained for a clade restricted to certain cyanobacterial and animal PADIs that excludes a fully supported outgroup clade containing fungal, actinobacterial and proteobacterial sequences (Figure S5). Full node support placed fungal PADIs in a clade with actinobacterial sequences to the exclusion of animal PADI sequences. This tree topology, whereby animal sequences have closer affinity to those in cyanobacteria than they have to other eukaryotic (fungal) sequences is surprising because it is inconsistent with the known species tree.

Phylogenetic methods, in particular phylogenetic trees built on single genes, are not infallible. It is therefore possible that the observed topology represents the failure of phylogenetic inference in the case of this individual gene, such that an artefact (e.g. model misspecification) might explain the affinity of the separate eukaryotic *PADIs* to different bacterial *PADI* types. This might be indicated for example by a changing tree topology under different models of rate variation. A common approach for an initial maximum likelihood analysis uses a best-fitting (maximum likelihood) fixed rate matrix of amino acid substitutions to produce the tree^56–58^. This can be confounded if there is evolutionary rate variation over different parts of the tree or deviation from typical protein substitution rates. In particular, attention has been drawn previously to heterotachous evolution, where the evolutionary substitution rate of a given site may change over time^59^. Heterotachy might be particularly plausible in the case of *PADI*, given the very divergent species represented in the analysis (animal, fungal, cyanobacterial, actinobacterial).

To validate the tree topology, and informed by the large tree, bacterial *PADI* sequences were subsampled to cover a broad representation of orthologous sequences for more computationally expensive phylogenetic analyses (selected to maximise diversity of bacterial *PADI* sequences as determined by varied representation of bitscore similarity to the animal *PADI* sequences). Three approaches were used to address possible heterotachous effects: a Bayesian approach that samples over different fixed empirical rate matrices^60^; a maximum likelihood approach using a mixture model of 20 different fixed amino acid rate matrices (C20)^61^; and a Bayesian approach that allows for infinite mixture model categories sampled from the alignment by making use of a Dirichlet process prior (CAT-GTR)^62^. These methods have been shown to be more robust to long-branch artefacts, and to saturated sequence artefacts more broadly, due to the more sophisticated treatment of across-site rate heterogeneity^63–66^.

All of the above analyses recovered a single topology that supports a clade of cyanobacterial and animal sequences to the exclusion of a clade of fungal and actinobacterial sequences (Figure 1a). Posterior probabilities or bootstrap values for this topology were high, approaching 100% for each of the four diverse methods (Figure 1b). The analysis was repeated using additional bootstrap algorithms, including the full non-parametric bootstrap, obtaining full support^67,68^. Topology constraint tests rejected a number of randomly generated trees to rule out possible specific biases in bootstrap resampling. Lastly, we generated a constrained tree for the expected model where eukaryotic PADIs are restricted to a monophyletic group. These alternative trees and constraint tree were all significantly rejected (p<0.0001) by multiple statistical tests including the AU-test^69–71^.

**Figure 1:**
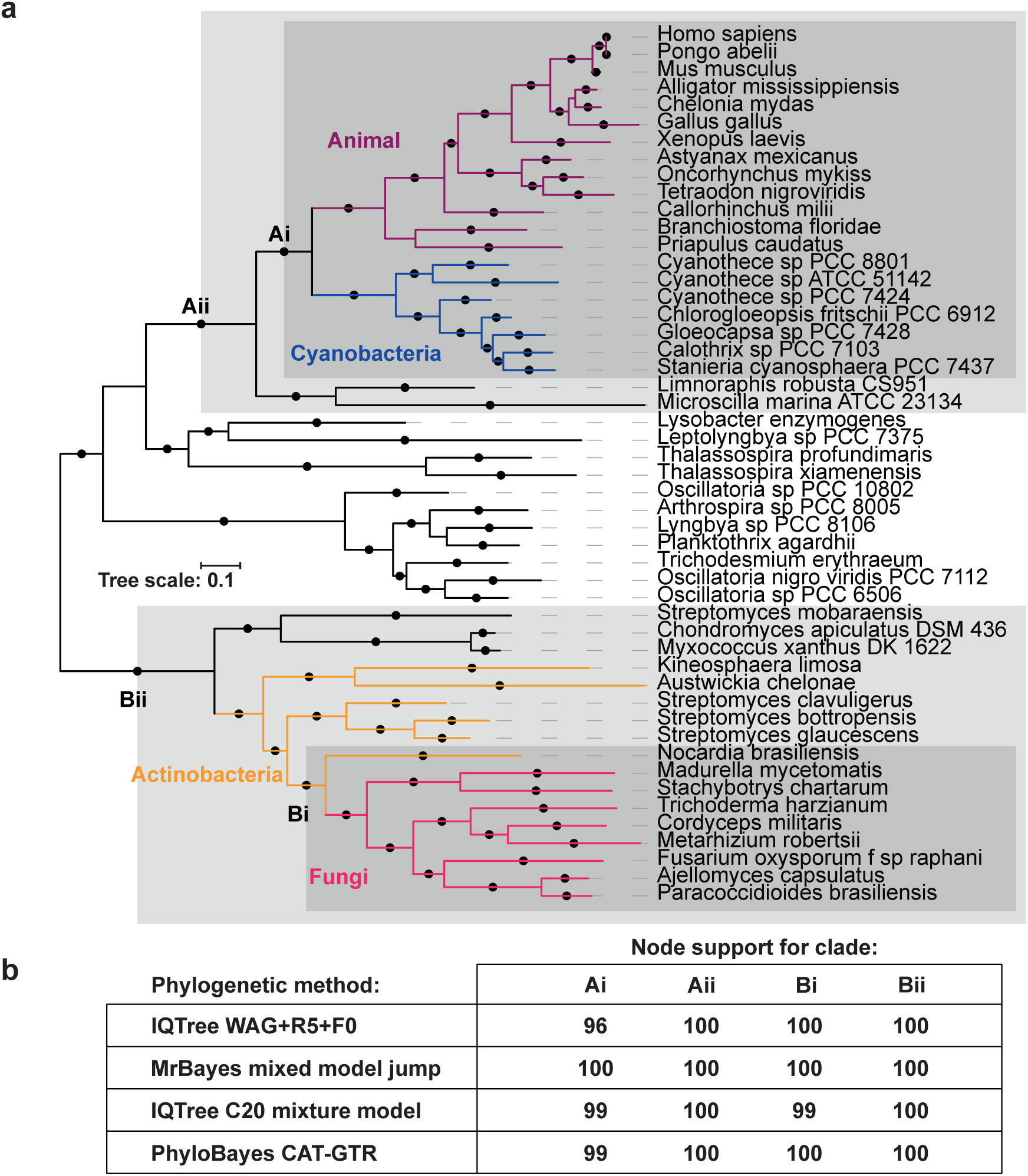
Phylogeny of the PADI sequence. **a)** Consensus topology for all phylogenetic methods with branch lengths from Bayesian phylogenetic inference with MrBayes. Solid circles indicate consensus node support of >95%. **b)** Node support for analysis with: 1) maximum likelihood inference using WAG fixed rate matrix, 2) MrBayes sampling across different fixed rate matrices, 3) IQTree using the C20 mixture model of rate matrices and 4) PhyloBayes using the CAT-GTR model of an infinite mixture model of rate matrices. Ultrafast bootstrap 2 values over 1000 replicates (for the Maximum likelihood methods) or posterior probabilities (for the Bayesian methods) are presented in the table for the nodes labelled in the tree critical to different evolutionary scenarios.

### Cyanobacterial and animal PADIs share unique synapomorphies

Although unlikely, some degree of stochastic evolution may potentially result in artefactual affinity of cyanobacterial *PADIs* with animal *PADIs* in the single gene tree. We therefore sought to identify features of the protein sequence that may independently validate the phylogenetic topology. A synapomorphy is any characteristic that is shared among multiple taxa but not with their ancestors. We identified three synapomorphies that are shared between animal and cyanobacterial PADI proteins but not with any actinobacterial or fungal PADIs (Figure 2).

**Figure 2:**
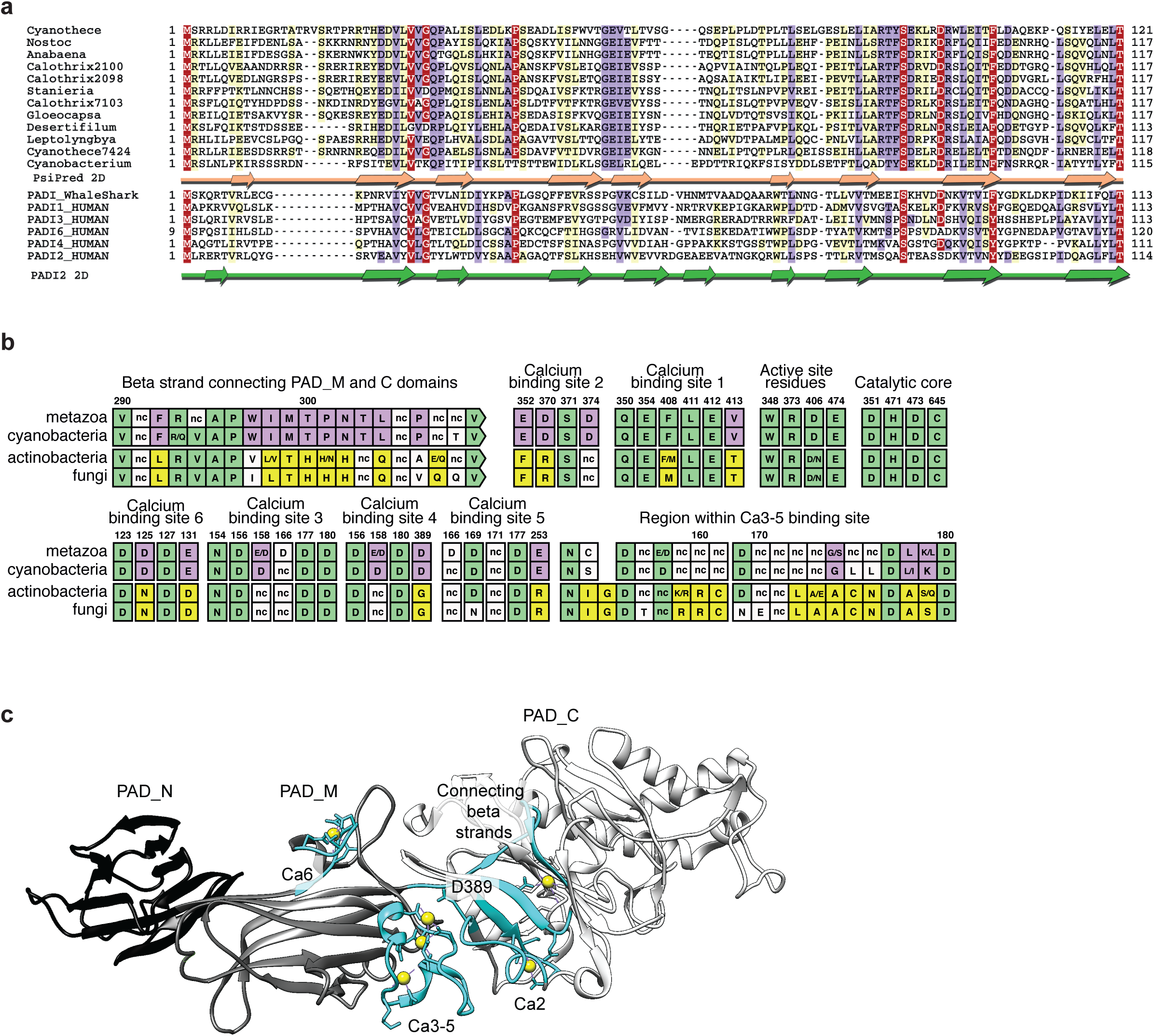
Synapomorphic features among PADI orthologues. **a)** Alignment of putative PAD_N domains from *SPM/NX* clade cyanobacterial PADI sequences with the PAD_N domain from human PADI paralogues and Rhincodon typus (whale shark). The colouring scheme indicates the average BLOSUM62 scores of each alignment column: red (>3.5), violet (between 3.5 and 2) and light yellow (between 2 and 0.5). Peach arrows shown below the cyanobacterial sequences indicate PsiPred predicted secondary structure (beta sheets). Green arrows (beta sheets) correspond to the known secondary structure of the PAD_N domain of human PADI2. **b)** Analysis of synapomorphic regions, representing six PADI sequences from each of metazoa, cyanobacteria, actinobacteria and fungi. Consensus sites across the six species are shown with standard single letter amino acid abbreviations. “nc” (non-nonserved) represents the absence of consensus conservation to one or two amino acids across the six species. The numbering given above the alignment and corresponds to the ungapped site of human PADI2 such that residues can be compared to Slade *et al*. Sites showing conservation across all four domains are coloured in green, sites showing synapomorphy between metazoa and cyanobacteria are coloured in purple, and sites showing synapomorphy between fungi and actinobacteria are coloured in yellow. **c)** Crystal structure of human PADI2 presented with PAD_N domain coloured in black, PAD_M domain in grey and PAD_C domain in white. Synapomorphic regions are coloured in cyan and calcium ions are shown as yellow spheres.

Firstly, we examined how the PADI protein domain architecture is distributed across orthologues using Pfam annotations, which are powered by HMMER searches^72^. As mentioned above, all metazoan PADIs possess the three PADI domains, PAD_N, PAD_M and PAD_C (Figure S1). The cyanobacterial PADIs closest to mammalian PADIs (from *SPM* and *NX* cyanobacteria) appear to possess two Pfam-annotated domains: a PAD_M domain and a PAD_C domain, but not a PAD_N domain. By contrast, other bacterial and fungal PADIs are only annotated with the PAD_C domain. To identify domains that might have been overlooked by Pfam, we carried out more sensitive profile-to-profile HMM searches^73,74^ (Figure S6a). We made a multiple sequence alignment firstly of cyanobacterial species contained in the monophyletic clade of metazoan sequences (Figure 1a, Clade Ai), and secondly of the remaining bacterial and fungal sequences (Figure 1a, sequences outside of Clade Aii). Regions corresponding to each of the PAD_N, PAD_M and PAD_C domains from human PADI2 were extracted and searched against a database of profiles of all domains contained in Pfam. This revealed that the bacterial and fungal sequences outside Clade Aii possess a divergent version of the PAD_M domain, but do not possess any PAD_N domain: the PAD_N region is completely absent from those fungal and bacterial orthologues, including cyanobacteria diverging earlier than *SPM/NX*. By contrast, the cyanobacterial homologues contained within Clade Ai (diverging after *SPM* and *NX* clades) possess all three domains including a degenerate metazoan PAD_N cupredoxin type domain (PAD_N domain: E-value<1×10^−7^). We then identified the cyanobacterial sequence that is predicted to assume the PAD_N secondary structure using PsiPred and aligned this with animal PAD_N sequences. The predicted cyanobacterial PAD_N sequence aligns well with the human PAD_N domain, as determined experimentally using PADI2 crystal structure data^28^ (Figure 2a), confirming that the Clade Ai cyanobacterial PADIs possess a degenerate PAD_N domain.

Secondly, we analysed representative fungal, actinobacterial, cyanobacterial, and metazoan PADI sequences for the conservation of calcium-binding and active site residues (Figure 2b). The allosteric binding of up to six calcium ions allows formation of the PADI2 active site cleft and is an absolute requirement for catalytic activity^28^. All catalytic residues and substrate binding residues are fully conserved among all PADI homologues (Figure 2b). In addition, calcium-binding sites 3 and 1 appear to be fully conserved, while calcium site 5 is also likely conserved. Ca6 is likely to be conserved functionally, as the substitution of D125 to N and E131 to D, which are present in both actinobacterial and fungal sequences, are expected to preserve ion binding. Intriguingly, however, calcium sites 2 and 4 appear to be exclusive to Clade Ai (late diverging cyanobacterial and metazoan) sequences. The fungal and actinobacterial sequences diverge from binding sites 2 and 4 to a different amino acid motif. Critically, only Clade Ai PADI sequences conserve the calcium switch residue D389 (residues: 369-389). In actinobacterial and fungal sequences this residue is substituted to Gly and therefore incompetent for metal coordination^28^ (Figure 2b). This indicates that the ordered, sequential calcium binding in the PAD_M domain, which is responsible for the allosteric communication between PAD_M and the catalytic PAD_C domain in human PADI2^28^ is likely to be conserved only in Clade Ai PADIs. As a result, a potentially different mode of calcium regulation operates in the fungal and actinobacterial PADIs.

Additionally, we find that fungal and actinobacterial sequences share features that are not present in the Clade Ai PADIs. This includes a conserved region within calcium binding sites 3-5 that is absent from the metazoan and cyanobacterial sequences (Figure 2b: amino acids 155-180, where differences conserved between fungal and actinobacterial sequences are highlighted in yellow). Also of interest is a highly conserved 10 amino acid beta sheet that connects the PAD_M and PAD_C domains (Figure 2b: amino acids 292-302). This region is conserved closely in fungal and actinobacterial sequences, but to a different 10 amino acid sequence containing a distinctive triple histidine motif (Figure 2b: amino acids 300-302).

The phylogenetic topology presented above is consistent whether built with or without the above synapomorphic sequence features and PAD_N domain (Figure S6b). As these synapomorphic features occur at the level of the amino acid sequence and at the level of a whole protein domain (Figure 2c), they are robust to convergent evolution, to differences in rate variation across the tree and to saturated sequence artefacts^75–78^. These features therefore provide strong additional support of the phylogenetic topology presented in Figure 1. It is inconceivable that blocks of sequence of up to ten amino acids were derived convergently and independently in actinobacterial and fungal PADIs. Thus these sequence features indicate a common ancestry of actinobacterial and fungal PADIs that is distinct from the ancestry of cyanobacterial and metazoan PADIs.

### The PADI sequence evolution between cyanobacteria and animals is anachronistically slow

The remarkably high similarity of Clade Ai cyanobacterial and animal PADIs prompted us to examine the rate of sequence change between them in more detail. We used a Bayesian phylogenetic approach to predict the divergence time between Ai Clade cyanobacterial and animal PADI sequences under a strict molecular clock model, using known fossil ages of metazoans as calibrations^79–81^. This prediction is expected to be approximately 3.35 billion years, in line with the age of the last common ancestor known to separate bacteria and eukarya^82^ (or greater, if evolutionary rates deviated in either lineage). Instead, our analysis yielded a far younger estimate of approximately 1 billion years (Figure 3a) for the age of their last common ancestor. Estimating divergence times using several relaxed clock models (UCLN, UCED, random local clocks^81^) increased the uncertainty in the estimate but, in all three cases, estimated an even more recent mean divergence time (Figure 3b). Under all approaches, the divergence times were not congruent with the geologically-defined divergence (*p* < 10^−8^) (Figure 3b). These divergence time estimates are therefore inconsistent with vertical descent of metazoan PADIs and are instead consistent with a horizontal acquisition event, which is more recent than the acquisition of the mitochondrion by eukarya. The divergence times predicted by these clock models are approximately dated at the time of divergence of the last common ancestor of PADI-harbouring metazoa.

**Figure 3:**
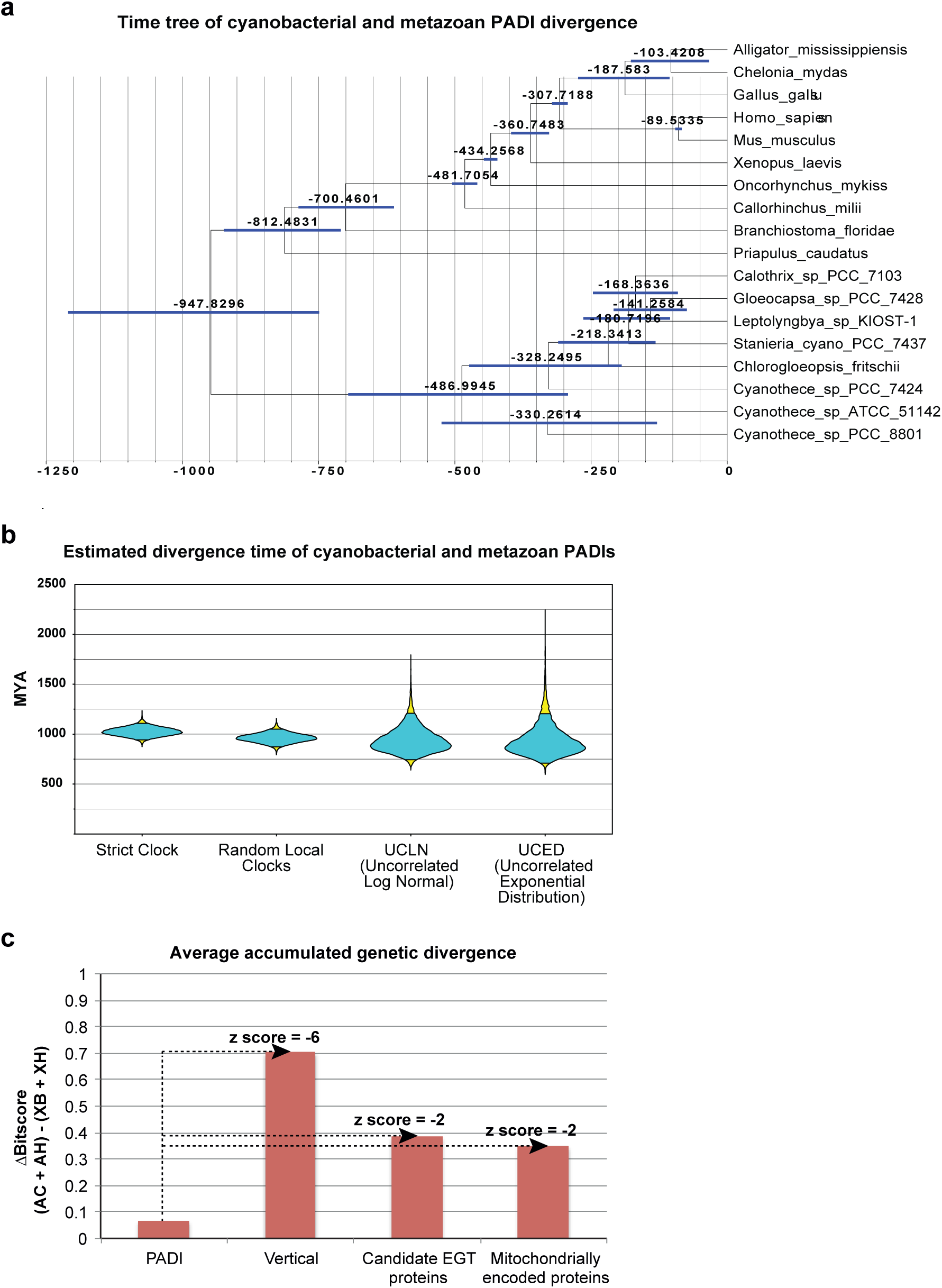
PADI sequence divergence analyses. **a,b)** Estimated divergence time of cyanobacteria and metazoa based on PADI sequences with respect to geologically defined constraints from the fossil record. **a)** Metazoan sequences and SPM/NX clade cyanobacterial sequences taken from the PATRIC database were used for Bayesian phylogenetic analysis in BEAST under the uncorrelated lognormal (UCLN) clock model, using a calibrated Yule model as the tree prior. Divergence times from the fossil record are used as normally distributed priors for six different nodes from metazoa. The marginal posterior distribution of the age of the root of the whole tree is used to estimate the divergence time; all nodes are labelled with the 95% credible interval for the marginal posterior distributions of the node ages. **b)** Estimated divergence times for running the analysis under different clock models (strict clock, random local clocks, UCLN and UCED), with the marginal posterior distribution of the age of the root of the whole tree given as a violin plot, where the 95% credible interval is given in cyan and the limits coloured in yellow. **c)** Analysis of the Accumulated Genetic Divergence (AGD) of PADIs compared to highly conserved proteins. The AGD was calculated for 26 vertically transferred proteins, 19 candidate EGT proteins, and 10 proteins encoded in the mitochondrial genome. The mean AGD for those groups is plotted against the AGD for PADI proteins.

Since the divergence of the PADI sequence between cyanobacteria and metazoa was unexpectedly low, we sought to understand it in the context of the genetic divergence between the species that bridge the closest PADI homologues. We therefore analysed a large number of the most conserved proteins in life to approximate a mean minimum extent of accumulated genetic divergence occurring between *Cyanothece sp. 8801* and *Branchiostoma belcheri* and compared this to the divergence of the PADI sequence between these two species (Figure S7). The distribution of calculated accumulated genetic divergence for 26 highly conserved proteins (ribosomal proteins, essential metabolic enzymes and chaperones) did not deviate significantly from a normal distribution (Figure S7). As a positive control for the hypothesised horizontal trajectory, we also analysed 19 proteins of likely endosymbiont gene transfer (EGT) origin and 10 proteins encoded in the mitochondrial genomes. Since mitochondrial and EGT-derived proteins were acquired more recently than the LUCA, the mean of the total accumulated sequence change for each of these proteins is expected to be much lower than that for vertically transferred genes (Figure S7). We therefore reasoned that they may mimic more closely the extent of accumulated genetic divergence that would be expected for an anciently horizontally transferred gene acquired more recently than the mitochondrion (as is hypothesised for the PADI gene). As expected, EGT and mitochondrially encoded proteins have an average accumulated genetic divergence that is significantly lower than that of vertically acquired proteins (Figure 3c). Specifically, the total accumulated genetic divergence for PADI sequences falls 6 standard deviations below that calculated for vertically transferred protein sequences, as assessed over the same timescale (Figure 3c). PADIs show less sequence change than all proteins individually analysed over this timescale and less even than ribosomal RNA. Indeed, they fall 2 standard deviations below the mean of EGT candidate genes or the mean of genes derived from the mitochondrial genome (Figure 3c). In a model of vertical descent, PADIs would therefore be under greater constraint than any other known sequence^83^.

### The cyanobacterial PADI protein is catalytically active

Considering the high degree of similarity between Clade Ai cyanobacterial and metazoan PADIs, including all necessary catalytic residues and calcium binding residues, we hypothesised that the ancestral cyanobacterial enzyme is likely to be catalytically active and calcium dependent. To test this, we prepared a recombinant version of the three-domain PADI from *Cyanothece sp. 8801* (here referred to as “cyanoPADI”) and assayed its catalytic activity alongside human PADI4. Analogously to the human enzyme, cyanoPADI can citrullinate multiple proteins in mouse cell lysates (Fig. 4a). In addition, cyanoPADI shows absolute dependence on calcium for activity. This demonstrates that the calcium-dependent regulation found in mammalian PADIs is also a feature of the ancestral cyanobacterial protein and suggests that the conserved calcium-binding sites, which were used in the evolutionary analysis as signifiers of synapomorphy, are functional (Fig. 2b and Fig. 4). Remarkably, and despite the absence of histones from bacteria, cyanoPADI catalyses citrullination of histone H3 (Fig. 4b), which is a known target of mammalian PADI4. The enzyme is additionally active at a physiologically relevant temperature for cyanobacteria (Fig. 4b). Thus cyanoPADI is a *bona fide* calcium-dependent peptidylarginine deiminase with sufficient similarity or promiscuity to catalyse citrullination of mammalian substrates.

**Figure 4:**
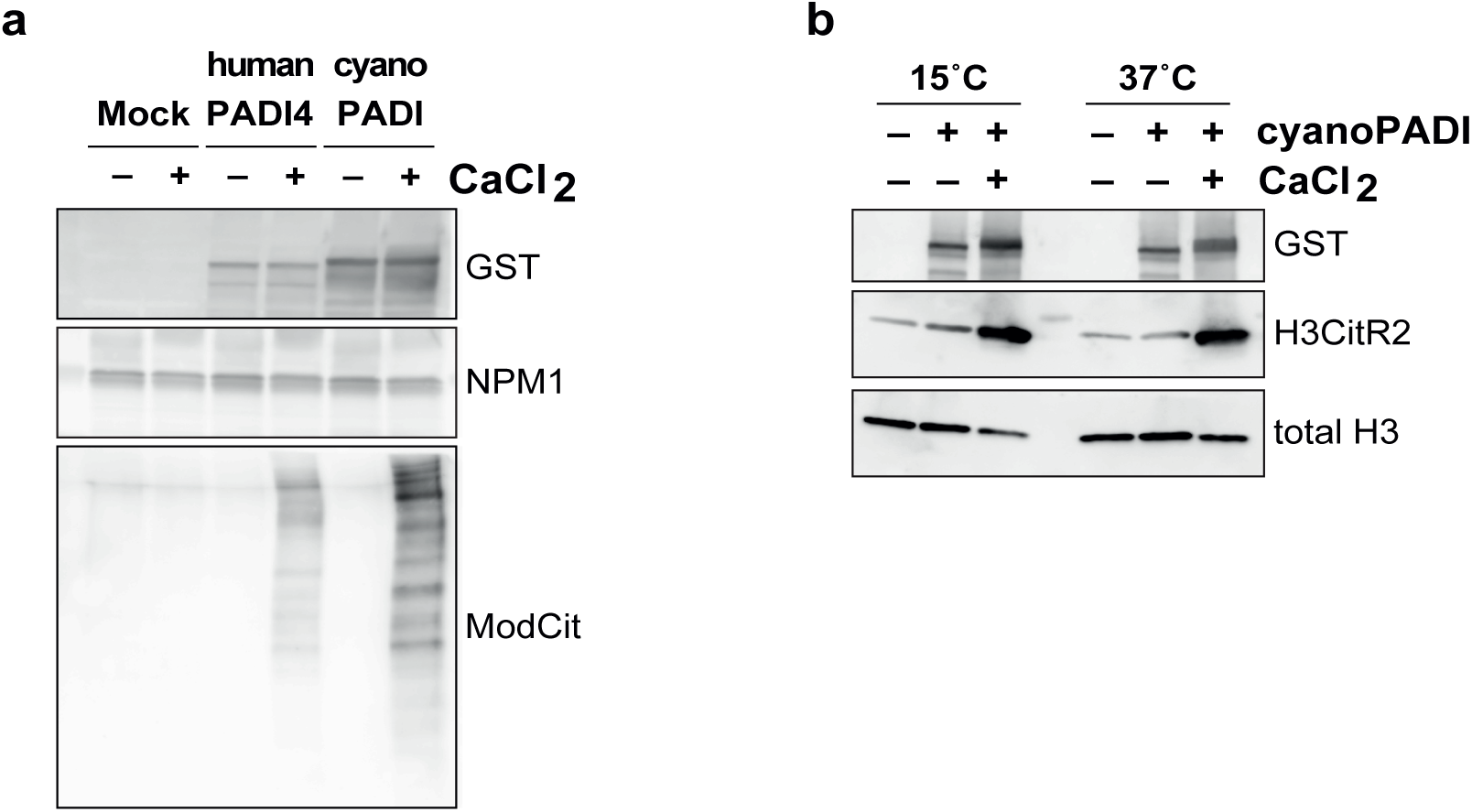
Biochemical analyses of the cyanobacterial PADI enzyme from Cyanothece sp. 8801 (cyanoPADI). Immunoblot analyses of citrullination assays using GST-His-tagged recombinant enzymes. **a)** Whole cell lysates from mouse embryonic stem cells were used as substrate and the presence of citrullination in a protein sequence-independent manner was assessed using the ModCit antibody. Nucleophosmin (NPM1) is used as a loading control. **b)** Recombinant human histone H3 was used as substrate and citrullination of arginine 2 was assessed. Total histone H3 is used as loading control.

## Discussion

It has been hypothesised that very few protein modification types existed in the LUCA and these have been diversified to give rise to the >200 PTMs known today^3^. We sought to map the evolutionary origin of citrullination, which is implicated in the regulation of an ever-increasing number of physiological and pathological processes. Our analyses of PADI homologues across life reveal the existence of two clearly discernible types of PADI homologues: one containing three structural domains and sharing functionally relevant sequence features and one containing two structural domains and divergent sequence features. The taxonomic distribution of these two types of homologues is highly unusual, in that three-domain PADIs are present in animal and late-diverging cyanobacteria, while two-domain PADIs are present in fungi and all other bacteria (Figures 1, 2, S6). This evidence can be reconciled with vertical evolutionary descent if the LUCA harboured two paralogous *PADI* genes which underwent widespread and mutually exclusive losses throughout evolution: firstly, the three-domain *PADI* present in late-diverging cyanobacteria and metazoa was lost from lineages leading to every other species in life; and secondly, the two-domain *PADI* present in fungi, actinobacteria and proteobacteria must be separately accounted for in independent gene losses in lineages leading to all other species. In lineages that harbour no *PADI*, the two paralogues must have been lost independently.

This highly unparsimonious scenario would be supported if rates of *PADI* sequence evolution across a species phylogeny were consistent with respect to geologically defined timings and with genes well known to have been inherited vertically from the LUCA. Our analyses of sequence divergence provide evidence to the contrary. In absolute terms, the similarity of cyanobacterial and branchiostomal PADIs to human PADIs is almost identical: 70.20% vs 70.90% respectively by pairwise amino acid similarity. However, a much greater amount of time has elapsed since the cyanobacterial and human genes have shared a last common ancestor than the genes from the other species pair (branchiostoma and humans). Even under assumptions of heterotachy, where rates of evolution may differ between different lineages, a minimal amount of nearly neutral genetic divergence nonetheless accumulates over evolutionary timescales in all lineages^83^. Under the assumption of vertical descent, the observed *PADI* sequence changes are anachronistically low even compared to the most highly conserved genomic sequences in life, including EGT candidates and mitochondrial genes.

The explanation for the observation of such little sequence change is more mundane under the assumption of horizontal transfer. A HGT event from late-diverging *SPM/NX* clade cyanobacteria to a last common ancestor within the animal lineage, although ancient, would have occurred much more recently than the LUCA and also likely more recently than the mitochondrion. HGT can therefore fully account for the phylogenetic distribution as well as the slow rates of evolution observed. The two lines of evidence are complementary and independent. The timing of transfer (neoproterozoic: 1000-542MYA) is consistent with the presence of marine nitrogen fixing cyanobacteria with specialised arginine catabolic pathways^84^, and with the emergence of metazoa in the cyanobacterial habitat^85–87^. A second HGT event, from actinobacterial species that are known to be fungal pathogens, most parsimoniously explains the existence of the two-domain fungal PADI (Clade Bi, Figures 1, S4 and Figure 2). This is consistent with the absence of a PADI gene either in eukaryotic species diverging before opisthokonts or in early diverging fungi such as yeast.

Closer examination of *PADI* phylogeny in bacteria provides additional support for HGT and confirms the directionality of horizontal transfer (Figure S8 and S9). Firstly, strong support is found for bacterial PADIs that form an outgroup to both the two-domain and three-domain PADI sequences (Figure S5 and S9). These bacterial outgroup sequences suggest that PADIs were not horizontally acquired by bacteria. Secondly, the fact that the metazoan-type three-domain PADI only emerges in the late-diverging *SPM* and *NX clades* of cyanobacteria, and the cyanobacterial PADI phylogeny mirrors the expected species tree^88^ (Figure S8), indicates that the three-domain PADI did not exist in the LUCA. The existence of cyanobacterial outgroup sequences, with a discernable origin within bacterial evolution, specifically implies the direction of HGT of the three-domain PADI was from cyanobacteria into metazoa and not in reverse (Figure S8).

All but one metazoan PADI sequence identified by our comprehensive searches in genomic and proteomic databases were found in deuterostomes – the exception being found in the *Priapulus caudatus* genome, a protostome. This suggests that the HGT took place either at the root of the deuterostomes, or possibly at the root of bilateria. Note that this part of the tree of life remains poorly resolved, with an extremely short branch between the bilaterian common ancestor and the deuterostomes^89^.

Biochemical analyses of the ancestral three-domain PADI (cyanoPADI) show that it is competent for catalysis (Figure 4), while a recent study has identified catalytically active PADI homologues in the thermotolerant fungi *Emericella dentata* and *Aspergillus nidulans*^90^. The discovery of catalytically active PADI orthologues in bacteria and fungi offers fertile ground for investigation of the roles of citrullination in these organisms.

Our finding that the cyanoPADI can citrullinate mammalian substrates (Figure 4) indicates that a novel catalytic capability was added to the regulatory repertoire of metazoan cells by HGT. The newly acquired regulatory function is likely to have enhanced biochemical diversity in animals. Fish genomes contain a single PADI gene, but duplications resulted in five tandem repeated paralogues in mammalian genomes^91^ (Figure S9). The fact that these duplicated genes were retained across many animal genomes suggests that they were unlikely to be functionally redundant. In the course of vertebrate evolution, citrullination was thus expanded in scope and adapted to a variety of cellular contexts, ranging from neutrophil extracellular trap release to stem cell potency, and from oligodendrocytes to bone marrow and keratinocytes^30^. The emerging physiological roles of the vertebrate PADIs, such as in the regulation of pluripotency and embryonic development^8,24,92,93^, or the newly described role of the fish PADI in tissue regeneration^94^, point to possible selective advantages conferred to metazoans by PADIs and offer a possible explanation for the fact that PADIs were retained so widely^95^. In a similar vein, it is interesting to consider our findings in light of the proposal that genes with a role antimicrobial defence are amenable to co-option by eukaryotic innate immune systems^46^. The extent to which the molecular mechanisms that regulate the human PADIs were also conserved from cyanobacteria or were newly co-opted in vertebrates remains an intriguing open question.

It is notable that no citrullination-reversing enzyme has been identified in any species to date. The evolutionary analysis of PADIs presented here adds extra complexity as to whether the reverse catalytic process might have also arisen or been propagated. It has been postulated that “toolkits” of PTM writer, eraser and reader enzymes may have evolved in a coordinated fashion and this has been studied formally in the context of protein phosphorylation^96^. In this context, the investigation into potential reverse catalysis for citrullination should be extended to include bacterial and fungal enzymes.

A related consideration is prompted by the known role of PADIs in autoimmunity. It has been proposed that the exogenous citrullinating activity of pPAD at sites of periodontal infection is an initiating event in the development of RA, by predisposing individuals with prior periodontal infection to the development of ACPAs^97^. It is therefore of note that pPAD and gADI genes are more widespread than previously thought (Figure S4) and that the PADIs described in this paper can be found in a number of human pathogens and in *Stachybotrys chlorohalonata* (black mold). A re-evaluation of the initiating events responsible for citrullination-specific breaks in immune tolerance may therefore be warranted.

This work reveals the remarkable evolutionary trajectory of the *PADI* family of human genes and uncovers the origin of a pleiotropic regulatory protein modification. In combination, the pieces of evidence presented above comprise a compelling case of ancient horizontal transfer of a bacterial gene into animals.

## Author contributions

T.F.M.C. and M.A.C. conceived the idea for the project and wrote the manuscript. T.F.M.C. and K.G. performed conservation, phylogenetic and time divergence analyses. T.F.M.C. performed domain architecture and structural analyses. L.S.P. performed structure-informed multiple sequence alignments. A.R.W. generated the vector for expression of recombinant cyanoPADI. G.G and T.F.M.C. performed protein expression and purification and carried out biochemical assays. C.D. and D.M advised on aspects of taxonomy and phylogeny. C.P.P. advised on aspects of structural and evolutionary biology. C.D and C.P.P. helped edit the manuscript.

## Acknowledgements

This work was funded by a Wellcome Trust and Royal Society Sir Henry Dale Fellowship and a MRC/University of Edinburgh Chancellor’s Fellowship to M.A.C. C.P.P. and L.S.-P. were funded by the Medical Research Council. D.M. and C.D. were funded by Swiss National Science Foundation Grant 183723. We thank M. Reijns for the gift of the pGEX-His plasmid, and G. Abrusán, G. Slodkowicz, M. Babu, N.D. Hastie and members of the Christophorou laboratory for critical discussions of the work.

## Materials and Methods

### Structural analyses

Structural homology searches were performed using the Dali server v3.1 with the extracted PAD_C domain used as query^98^. Superposition of known structures was performed in Chimera^99^ using the MatchMaker tool^100^. Briefly, the two structures (PDB: 4n2c and 1xkn) were aligned for the best-aligning pair of chains using the Needleman-Wunsch algorithm and BLOSUM62 matrix. A secondary structure score of 30% was included. The superposition was iterated by pruning long atom pairs such that no pair exceeds 2.0 angstroms.

### Identification of PADI orthologues

A graph-based unsupervised clustering algorithm used by the EggNOG database was used to infer PADI orthologous groups from 2031 genomes across the tree of life (ENOG410ZKF3: 217 proteins from 74 species)^55^. Phylogenetic reconstruction for the identified PADI orthologues was performed within EggNOG as implemented within the ETE3 suite (eggnog41) and described at http://eggnogdb.embl.de/#/app/methods^55,101^. In addition, a list of proteins with significant similarity (E-value < 1×10^−3^) to the metazoan PAD_C domain from human PADI2 were collected using HMMER searches against *Reference Proteomes* and *UniProtKB* databases^102^. Additional more sensitive sequence searches and iterative searches were performed using tblastn and psiblast against nr/nt; jackhammer against reference proteomes and *UniProtKB*; and hhpred against Pfam-A, COG_KOG and PDB_mmCIF70^73,103,104^. To verify the exhaustive nature of our search for PADI homologues we employed two state of the art remote homology detection tools. The first, HHBlits, is used to search databases of hidden markov models (hmm) generated with clustered proteomic datasets with a query hmm and is available in the HHSuite^105,106^. The second tool, Hmmsearch, is included in the Hmmer suite^107^ and used to search proteomic datasets with an hmm. A PADI alignment was generated from our initial dataset of known homologues for use with both tools using clustal omega on default parameters for 3 iterations to generate our query hmm. HHBlits was used to search the Uniclust30 database. Its construction and contents are detailed on the mmseqs website^108^ (Uniclust. [cited 5 May 2020]. Available: https://uniclust.mmseqs.com/). Hmmsearch was used to search the NCBI NR protein database. Its contents and construction are detailed on the NCBI web page (Download-NCBI. [cited 5 May 2020]. Available: https://www.ncbi.nlm.nih.gov/home/download/). The HHBlits search results were filtered with a cutoff of 90% probability and 50 amino acids. No additional hits were found to sequences in clades that were not in the starting dataset.

### Analysis of spurious viral and archaeal hits

The HMMsearch results were filtered with an E-value cutoff of 10^−10^ and 50 amino acids. Two sequences attributed to unexpected clades were found: refseq identifiers AXN91134.1 and RCV64870.1 which are found in *Namao virus* and *Methanophagales archaeon*, respectively. The two anomalous sequences are the only representatives of the *PADI* family within their taxonomic kingdoms and this extremely sparse distribution of these sequences would either imply many independent gene loss events or an extremely recent horizontal transfer event of *PADI* to these clades if the genes are in fact correctly attributed to their genome. To verify the validity of the attribution of these sequences to their respective genomes, we calculated a phylogeny including by aligning a subset of high confidence *PADI* sequences with the two putative homologues. The alignment was then used with IQTree^109^ on default parameters and automatic selection of the appropriate model to generate a phylogeny. The resulting tree was visualized with figtree^110^ (Figure S2). The placement of the two sequences in the phylogeny does not agree well with a plausible evolutionary scenario considering their taxonomic origin; rather their placement suggests that the allegedly *Methanophagales archaeon* sequence is in fact a cyanobacterial sequence, and that the allegedly *Namao virus* sequence, is in fact a fish sequence. These hypotheses are corroborate by the origin of the samples used to obtain them: metagenomic isolates in the case of *Methanophagales archaeon*, and infected tissue samples taken from fish in the case of *Namao virus*. In both cases, the sample were susceptible to gene misattribution due to incorrect binning^111^ or contamination. To further probe the association of these genes to their respective genomes and discount the possibility of the *PADI* genes belonging to a transferred genomic segment, we used the K-mer spectra of the genomes to study the possibility of horizontal transfer events. These analytics are regularly used to find transferred regions in prokaryotic genomes^112^. Normalized K-mer spectra for DNA sequences were generated by counting occurrences of all K-mers and normalizing by the total amount of words counted to give a unit vector. The results presented in Figure S3 were derived using 4-mers. To detect possible horizontally transferred genomic regions, an average spectrum for the entire genome was calculated. A spectrum was then calculated for a sliding window of 1 kb using 500bp steps and subtracted from the genomic average at each window position. The absolute value of the difference between the genomic average and window spectra is represented over the entire genome. The code for running these kmer-based analyses is available at https://github.com/DessimozLab/Pad1.

### Phylogenetic methods

For all phylogenetic trees, branch support information was visualised and figures produced using FigTree v1.4.3 and iTOL^113^. Amino acid sequences for PADI homologues were obtained from UniProtKB, NCBI and Pathosystems Resource Integration Center (PATRIC) databases using HMMER and BLAST searches^72,114,115^. PADI2 was used for species with multiple PADI paralogues, as it closest resembles the PADI gene in metazoa with one PADI (such as fish^116^), and with the PADI2 from metazoan species with three PADIs such as birds or reptiles (Figure S7).

### Phylogenetic analysis of other citrullinating enzymes

Sequences of the arginine deiminase from *Giardia lamblia* (gADI^33^) and the porphyromonas-type peptidylarginine deiminase from *Porphyromonas gingivalis* (pPAD^117^) were used as a seed for Hidden Markov Model (HMM) searches of reference proteomes to identify sequences from other species of similar length and most significant similarity^72,102^. These were aligned with 25 representative PADI sequences using MAFFT L-ins-I^118^ and singly aligning columns were removed. IQTree was used to produce a maximum likelihood phylogenetic tree^109,119^. The LG empirical rate matrix with 8 categories of rate variation under the FreeRate model (LG +R8) was used, as determined by ModelFinder^58,120^ according to the corrected Akaike Information Criterion. The Ultrafast Bootstrap 2 with 1000 replicates^67^, Shimodaira-Hasegawa (SH)-like approximate likelihood-ratio test (aLRT) with 1000 replicates^121–123^, and aBayes parametric tests^124^ were used to assess node support.

### Phylogenetic analysis of PADI orthologues

All putative bacterial PADI sequences in the PATRIC database were obtained from BLAST searches^72,114,115^. In addition, sequences from metazoa were subsampled to maximise the inclusion of different lineages. The human PADI2 sequence was searched against *UniProtKB* to subsample sequences from 35 fungal species that represent the broadest distribution of affinity with PADI2 as determined by HMMER bitscore^102^. This was done to provide representation from the maximum sequence diversity of fungal PADIs. Fragment sequences were excluded. The collected sequences were aligned using MAFFT L-ins-I and singly aligning columns were removed^118^. IQTree was used to produce a maximum likelihood phylogenetic tree^109,119^. The WAG empirical rate matrix with 10 categories of rate variation under the FreeRate model with base frequencies counted from the alignment (WAG +R10 +F) was used as determined by ModelFinder according to the corrected Akaike Information Criterion^57,58^. Ultrafast Bootstrap 2 with 1000 replicates, SH-like aLRT with 1000 replicates, and aBayes parametric tests were used to assess node support^67,121–124^. The tree is shown rooted at the midpoint with solid circles indicating consensus node support of >95%. A number of critical nodes for testing different evolutionary hypotheses were labelled in full as they are mentioned specifically in the text or referred to in other analyses.

### Phylogenetic analysis of subsampled PADI orthologues for topology testing

Bacterial PADI sequences were subsampled to maximise sequence diversity. Both the closest and the most distant bacterial homologues with respect to the metazoan sequence were retained to allow for the broadest distribution of protein sequences (to a total of 50 sequences). Firstly, a maximum likelihood phylogenetic tree was produced using IQTree as above using the best empirical rate matrix according to ModelFinder (WAG + R5 +F0). This reproduced the topology obtained with the full tree. Three further analyses were performed:

1. Maximum likelihood phylogenetic analysis was performed using IQTree under the C20 empirical profile mixture model of evolutionary rate matrices using CIPRES Gateway on XSEDE^61^.
2. Bayesian phylogenetic inference was performed using MrBayes v3.2.6 x64 using CIPRES Gateway on XSEDE with mixed model MCMC jumping across different fixed empirical rate matrices and 5 different gamma distributed rate categories^60^. Analysis was performed with 4 runs each of 1000000 chains. The average standard deviation of split frequencies was observed to be <0.005, parameters all had an effective sample size (ESS) > 500 and potential scale reduction factor (PSRF) of 1.000 (to 3 significant figures). The summary tree was generated with a burn-in of 25% over the runs. Posterior probability was used for node support– i.e. where posterior probability was 100, the topology was congruent in every tree sampled by the Markov chain Monte Carlo (MCMC) after burn-in. The *aminoacid model* prior was set as ‘mixed’ such that the MCMC jumps across different models i.e. mixture of models with fixed rate matrices. Poisson, Jones, Dayhoff, Mtrev, Mtmam, Wag, Rtrev, Cprev, Vt and Blosum models were used and all have equal prior probability. The WAG model had posterior probability of 1.000, and standard deviation <0.0001 – and was exclusively sampled from the posterior^57^. This is consistent with the WAG model being identified as the best empirical matrix identified according to ModelFinder and the corrected Akaike information criterion from the maximum likelihood analysis in IQTree.
3. Bayesian phylogenetic inference was performed using PhyloBayes under the CAT-GTR model^62,125^. This is an infinite mixture model of rate matrices making use of a Dirichlet process prior. The alignment contains 1100 aligned positions and 50 taxa. 8 chains were performed in parallel for 24 hours such that more than 20000 cycles were achieved as recommended in the PhyloBayes manual using the MRC IGMM and University of Edinburgh computing cluster Eddie3. Readpb, bpcomp, tracecomp tools in PhyloBayes and Tracer software were then used to analyse runs. Posterior consensus trees were generated for each run and were reproducible across the eight different runs. The trace plots for independent runs were also analysed to assess for apparent stationarity aiming for an ESS of at least 100. Maxdiff was observed to be < 0.1 (maxdiff= 0.06209, meandiff= 0.00330).

Tree topologies were congruent across the four different methods (Figure 1b). Additional topology testing was performed in PAUP*4.0a163 and in IQTree. 100 random trees were generated along with the maximum likelihood constrained tree where fungal and metazoan sequences were constrained to be monophyletic. The SH-test, approximately unbiased (AU) test and expected likelihood weight (ELW) tests were performed and all other alternative trees, including the constraint tree were rejected (p<0.0001)^69–71,122^.

### Phylogenetic analysis excluding synapomorphic regions

Phylogenetic analyses from Figure 1 were repeated using an alignment with the PAD_N domain removed and with an alignment in which both the PAD_N domain and regions of synapomorphy (Figure S4) were removed. Maximum likelihood analysis using IQTree with ModelFinder used to select the best performing fixed empirical rate matrix (WAG + R5 +F0) as above^58,67,109,119^. Topologies were congruent with analysis of the whole alignment and node support values (Ultrafast Bootstrap 2) for the clades labelled in Figure 1 are provided in Figure S4b.

### PADI Domain annotation

To identify putative locations for the three PAD domains within PADI homologue sequences from bacteria and fungi, each target PADI sequence was aligned to five metazoan sequences using TCoffee^126^. Putative domain sequence regions were then used as a target query for HMMER or HHPred searches^72,127^. HMMER searches were made against the *UniProtKB* database and HHPred searches were made, firstly against a database of HMM profiles of protein domains in the Protein Data Bank (PDB_mmCIF_4_Aug) and secondly, against a database of profiles from Pfam (Pfam-A_v31.0)^104^. Once individual sequences were identified as possessing a specific domain architecture, multiple sequence alignments of groups of sequences with common putative domain architecture were made and this was used as a query for each type of search.

For the reported E-values in Figure S4a the following method was used. All sequences from the highlighted clade in the phylogenetic tree were aligned using TCoffee, PAD_C, PAD_M and PAD_N domains extracted from the cyanobacterial sequences, and secondly PAD_C and PAD_M domains from the clade containing a mixture of bacterial and fungal sequences. This alignment was used as a seed for searches with HHPred against a database of profiles made of the entire human proteome, and against a database of profiles of Pfam domains (Pfam-A_v31.0). HHPred searches were performed using the MPI Bioinformatics Toolkit of the Max Planck Institute for Developmental Biology, Tübingen, Germany^74,104^.

### Multiple sequence alignment of PAD_N domain

Amino acid sequences were aligned using MUSCLE or TCoffee algorithms^126,128^ and visualised using Jalview^129^. Putative PAD_N domains from *SPM*/*NX* clade cyanobacterial PADI sequences were identified using HHPred showing significant statistical evidence for affinity (E-value: 2.5×10^−5^)^74,104^. These were aligned with the PAD_N domain from human PADI paralogues and *Rhincodon typus* (whale shark). The alignment was presented with the program Belvu using a colouring scheme indicating the average BLOSUM62 scores (which are correlated with amino acid conservation) of each alignment column^130^, as represented in Figure 2a. PsiPred^131^ was used to predict secondary structure for the cyanobacterial PAD_N domains (beta sheets). The secondary structure of the PAD_N domain of human PADI2 was identified from the crystal structure (PDB: 4n2a)^28^.

### Synapomorphy analysis of PADI calcium binding sites

Representative fungal, actinobacterial, cyanobacterial, and metazoan PADI sequences were analysed for the conservation of all of the calcium-binding sites (a minimum of three residues coordinate each calcium binding site) and for other critical residues contained at the active site (Figure 2). PADIs from the following species were used: 1) metazoan PADIs from *Homo sapiens, Xenopus laevis, Oncorhynchus mykiss, Callorhinchus milii, Branchiostoma floridae, Priapulus caudatus*; 2) cyanobacterial PADIs from *Cyanothece sp. 8801, Stanieria cyanosphaera, Chlorogloeopsis fritschii PCC 6912, Crocosphaera subtropica, Aphanothece sacrum, Cyanothece sp. 7424;* 3) fungal PADIs from *Fusarium sp. FOSC 3-a, Periconia macrospinosa, Paracoccidioides lutzii, Blastomyces parvus, Ajellomyces capsulatus, Emmonsia crescens* and; 4) actinobacterial PADIs from *Streptomyces silvensis, Alteromonas lipolytica, Streptomyces sp. 3214*.*6, Erythrobacter xanthus, Kibdelosporangium aridum, Nocardia brasiliensis ATCC 700358*. Sequences were aligned using MAFFT L-ins-I and compared to functionally annotated regions from *Slade et al*. 2015 and from crystal structures^28,34,118^.

### Sequence divergence analyses

BEAST v2.4.8 was used to produce a time tree of the clade of subsampled metazoan PADIs and the full clade of closest *SPM*/*NX* cyanobacteria contained within the PATRIC database using the GTR model with 4 gamma distributed rate categories^79,88,132^. The following metazoan species were used: *Homo sapiens* (HS), *Mus musculus* (MM), *Alligator mississippiensis* (AM), *Chelonia mydas* (CM), *Gallus gallus* (GG), *Xenopus laevis* (XL), *Oncorhynchus mykiss* (OM), *Callorhinchus milii* (CM), *Branchiostoma floridae* (BF), *Priapulus caudatus* (PC). To calibrate nodes on the tree, node times were set as the following normally distributed priors: mean 797.0, sigma 72.5 (clade of HS, MM, AM, CM, GG, XL, OM, CM, BF, PC); mean 692.5, sigma 57.5 (clade of HS, MM, AM, CM, GG, XL, OM, CM, BF); mean 473.5, sigma 14.0 (clade of HS, MM, AM, CM, GG, XL, OM, CM); mean 435.0, sigma 6.5 (clade of HS, MM, AM, CM, GG, XL, OM); mean 311.0, sigma 7.5 (clade of HS, MM, AM, CM, GG); mean 89.5, sigma 3.0 (clade of HS, MM). Metazoan divergence times from the fossil record were obtained from timetree.org with bounds on the distributions chosen to span the range of times reported in the literature centered on the median value^133^. The calibrated Yule model was used as the tree prior. XML files were generated in BEAUti and the MCMC analysis was run using BEAST2 on the CIPRES Gateway on XSEDE. An initial MCMC run of 5,000,000 chains was run for each clock model^80,81^. Then full analysis was performed with two independent runs of 10,000,000 chains for the strict clock, uncorrelated lognormal (UCLN) and uncorrelated exponentially distributed (UCED) clock models and three independent runs of 10,000,000 chains for the random local clocks model. All models were additionally run under the tree prior (i.e. in the absence of sequence data) and assessed. Analysis of parameters was performed in Tracer to assess apparent stationarity for the different tree parameters and for acceptable ESS values and congruence was assessed across the independent runs. The predicted divergence time of the metazoan and cyanobacterial clades was given by the marginal posterior distribution of the age of the root of the whole tree. This is given by the TreeHeight parameter. The 95% confidence interval of the TreeHeight parameter for all runs under each of the clock models did not exceed 1300. This data was plotted as a violin plot showing 95% confidence intervals (Figure 3b).

### Accumulated genetic divergence analysis relative to other proteins

Bitscore density is calculated by taking the bitscore of a query sequence to the target sequence produced by HMMER and dividing by the bitscore of the query sequence to itself (longer sequences have higher bitscores), which gives a value between 0 and 1^72^. The bitscore densities of the similarity of 1) the cyanobacterial homologue to the human sequence: ΔbitscoreD_Cy-Hu_ (AC+AH) and 2) of the branchiostomal homologue to the human sequence: ΔbitscoreD_Br-Hu_ (XB+XH) were both calculated (Figure S5). A measure of the total accumulated genetic divergence between late-diverging cyanobacteria (*Cyanothece spp*.) and the last common ancestor of *Branchiostoma spp*. and *Homo sapiens* was then calculated by subtracting ΔbitscoreD_Br-Hu_ from the ΔbitscoreD_Cy-Hu_. This accumulated genetic divergence (AGD) value was calculated for: (1) 26 ribosomal proteins (uS2, uS3, uS4, uS5, uS7, uS8, uS9, uS10, uS11, uS12, uS13, uS17, uS19, uL1, uL2, uL3, uL4, uL5, uL6, uL11, uL13, uL14, uL15, uL22, uL23, uL24), (2) 19 sequences whose proteins are mitochondrially located so are reasonable EGT candidates from the mitochondrion (OTC, ASS1, ARLY, CPS1, PGK, ENO, GAPDH, PK, NAXE, G6PD, RPIA, FUMH, SDHB, SDHA, CS, MDHM, DLAT, DLDH, ACLY)^134^, and (3) all 10 proteins still encoded in the mitochondrial genome (MT-ATP6, MT-CO1, MT-CO2, MT-CO3, MT-CYB, MT-1, MT-2, MT-3, MT-4, MT-5). It is notable that by definition, only very highly conserved proteins have an AGD that can be calculated in this extreme example between the last common ancestor of late diverging cyanobacteria and humans: if a protein has diverged substantially then the similarity of the human homologue to the cyanobacterial will not be discernible and no bitscore can be calculated. AGD values of proteins in each category were then tested for deviation from a normal distribution using the Shapiro Wilk test (*W* = *b*^*2*^ / *SS*)^135^. As the calculated p-value > 0.05, the null hypothesis was retained and the data treated as being normally distributed. Kurtosis and skew were also within the range of the normal distribution. The AGD for PADI proteins (AGD_PADI proteins_ = 0.066) was then compared to the mean AGD of each category of control proteins (e.g. AGD_ribosomal proteins_ = 0.70) and z-scores were calculated.

To compare the extend of divergence relative to ribosomal RNA (rRNA), nucleotide sequences for rRNA were obtained from the SILVA database^136^. Nucleotide sequences for PADIs were obtained from NCBI and exons extracted. Comparisons were made with EMBOSS Needle using the Needleman-Wunsch global alignment algorithm^137^ (gap open: 10, gap extend: 0.5).

### Preparation of recombinant proteins

PADI gene sequences were obtained from NCBI and synthesised by Thermo GeneArt with flanking EcoRI (at the 5’ end) and XhoI (at the 3’ end) restriction sites. *Cyanothece sp. 8801* PADI and human PADI4 sequences were subcloned into a modified pGEX vector, which included an additional 10X His tag immediately N-terminal of the enzyme sequence (generous gift from Dr Martin Reijns, MRC Human Genetics Unit), by InFusion cloning. GST-His-PADI4 and GST-His-cyanoPADI were expressed in BL21 (DE3) in 2TY cultures. Cells were grown (37 °C; 180 rpm) to an OD600 of 0.6 and induced overnight at 18 °C with 0.5 mM β-D-1-thiogalactopyranoside (IPTG). Bacterial pellets were harvested by centrifugation (8,000 xg; 10 min) and frozen at −80°C. Cell pellets were resuspended in 50 mM Tris pH 7.5, 500 mM NaCl, 20 mM imidazole, 5% Glycerol, 1 mM DTT (1 g dry cell mass in 4 mL), 1X EDTA-free protease inhibitors (Roche), 5 mM MgCl_2_ and 10 units benzonase at 4°C with stirring. Cells were lysed on ice by sonication (7x 45 sec, with 45 sec breaks) and the lysate was cleared by centrifugation (20,000 xg; 20 min). Supernatant was sterile filtered (0.2 µm) before loading by Superloop onto a 5 mL HisTrap column, which was pre-equilibrated with binding buffer. Proteins were purified using an AKTA FPLC system (GE Healthcare). The column was washed with 50 mM Tris pH 7.5, 500 mM NaCl, 25 mM imidazole, 5% Glycerol, 1 mM DTT and the recombinant proteins were eluted with 50 mM TRIS pH 7.5, 500 mM NaCl, 250 mM imidazole, 5% Glycerol, 1 mM DTT. The purified sample was concentrated using Vivaspin MWCO filters into 50 mM HEPES pH 7.5, 150 mM NaCl, 5 mM DTT, 5% (v/v) glycerol, and the concentration determined using Nanodrop.

### Citrullination activity assays

#### Using mouse cell lysates

E14 mouse embryonic stem (ES) cells were cultured in GMEM supplemented with 10% fetal calf serum (FCS), 0.1 mM non-essential amino acids, 2 mM L-glutamine, 1 mM sodium pyruvate, 0.1 mM beta-mercaptoethanol and 10^6^ units/L leukaemia inhibitory factor (LIF) (ESGRO, Millipore) and grown on a six well plate until 70% confluent. Cells were harvested in 0.1% NP-40, 20 mM Tris pH 7.6, 1X EDTA-free protease inhibitors, 5 mM DTT, after two washes in PBS (one in PBS containing 2 mM EDTA and one in plain PBS). To shear chromatin and clarify lysates, benzonase and 2 mM MgCl_2_ were added and samples were rotated at 4°C for 30 mins, sheared by passing through a 25G needle and centrifuged at 13000 rpm for 5 mins. Citrullination activity assays were performed with 500 nM recombinant enzyme in 50 mM HEPES pH 7.5, 150 mM NaCl, 5 mM DTT, 5% (v/v) glycerol, either in the presence of 5mM CaCl_2_ or water. Reactions were incubated for 30 mins at 37°C and quenched by boiling at 95°C for 5 mins. Samples were stored at −80°C before immunoblotting.

#### Using recombinant histone H3 substrate

Reactions were performed in 50 mM HEPES, 137 mM NaCl, 5 mM DTT with 1.5 µM recombinant H3 (New England Biolabs), vehicle or 500 nM recombinant enzyme, and in the presence of either 5 mM CaCl_2_ or water. Reactions incubated for 30 mins at 15°C or 37°C and quenched by boiling at 95°C for 5 mins before immunoblotting.

#### Immunoblotting

Proteins were separated by SDS-PAGE and transferred to nitrocellulose membrane using wet transfer. Membranes were blocked in 5% BSA in TBS containing 0.1% Tween-20 for 1 h at room temperature. Proteins were detected using primary antibodies against anti-H3 (Abcam ab10799, 1:1000), anti-H3CitR2 (Abcam ab176843, 1:2000), anti-NPM1 (Abcam ab37659, 1:200) and anti-GST (Abcam ab19256, 1:1000) overnight at 4°C and in secondary antibody at 1:5000 for 1 h at room temperature. Membranes were incubated in Pierce ECL reagent and imaged using ImageQuant LAS 4000 (GE). Citrulline-containing proteins were modified on the membrane and detected using the anti-modified citrulline detection kit (Millipore, 17-347) according to manufacturer’s instructions.

**Figure S1:**
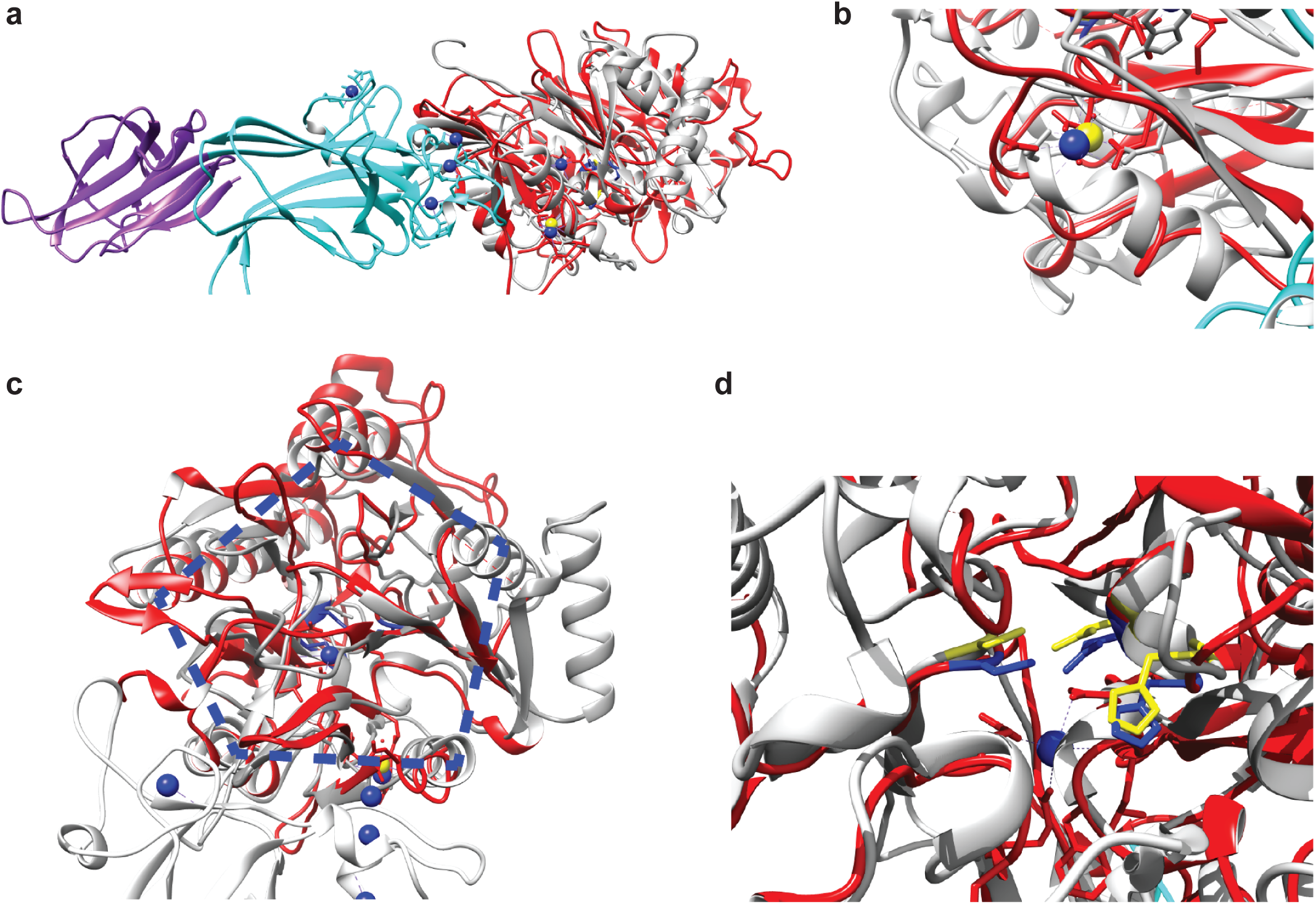
The active site of human PADI2 adopts an ancient configuration. PADI2 structure (PDB: 4n2c) in purple (PAD_N domain), cyan (PAD_M domain) and red (PAD_C domain) was superimposed with the structure of Agmatine deiminase from Chlorobium tepidum (ctAgD) in grey (PDB: 1xkn). **a)** Full structure of PADI2 shows that Agmatine deminase (grey) superimposes onto the PAD_C pfam domain (red). **b)** Detail of the conserved metal ion binding site between the two enzymes (Calcium binding site 2 in PADI2, blue sphere, and Na+ site in ctAgD, yellow sphere). **c)** Detail of the PAD_C domain shows the conservation of the overall pentein fold (blue dotted line shows five-fold rotational symmetry). **d)** Active site configuration and positioning is conserved between PADI2 (blue: His-Asp-Cys triad) and ctAgD (yellow: His-Asp-Cys triad).

**Figure S2:**
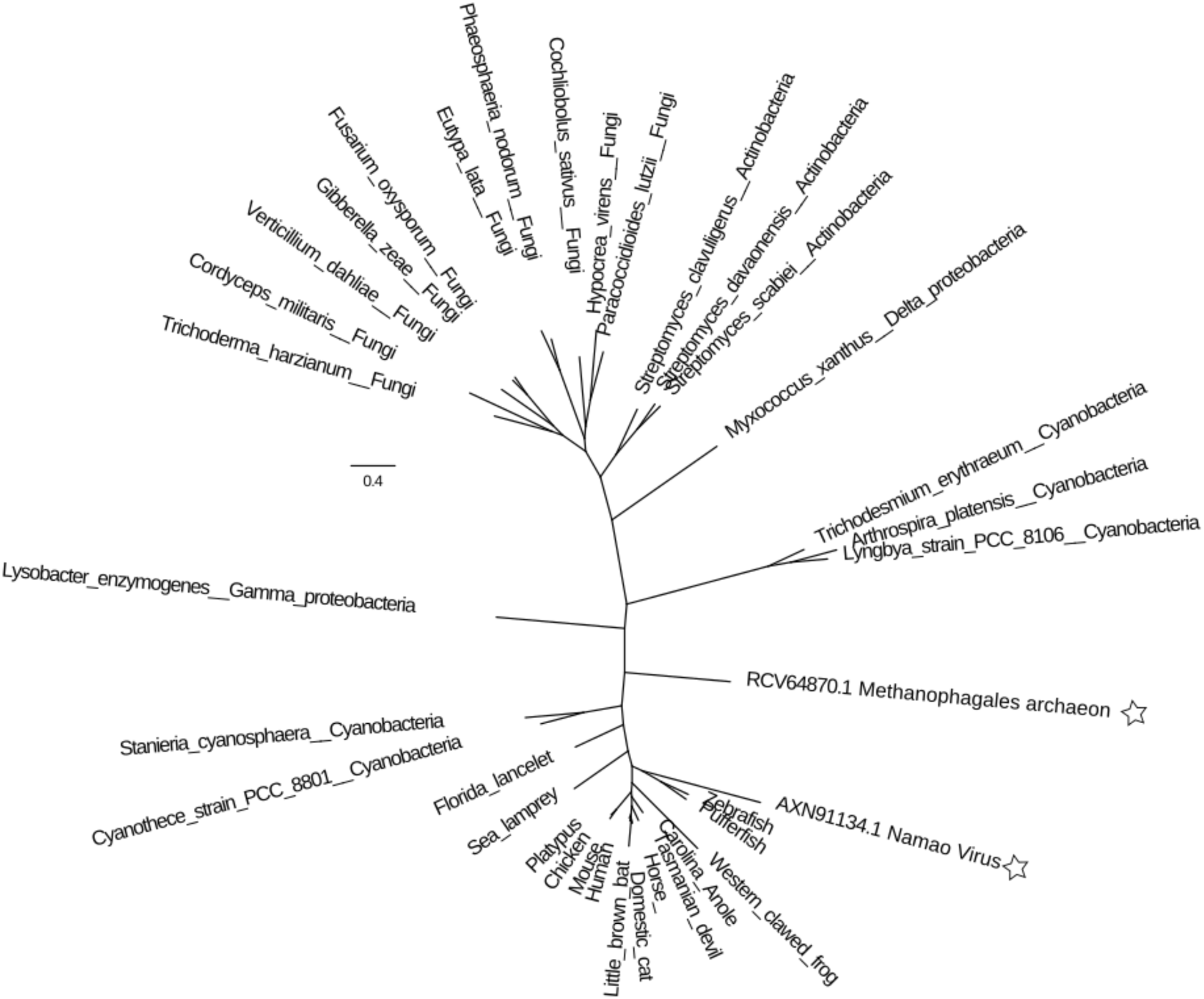
Phylogeny of two spurious putative *PADI* sequences from unexpected clades within the core set of *PADI* sequences. The putative *PADI* homologues from *Namao virus* and *Methanophagales archeon* are indicated with a star icon. The phylogeny of *PADI* homologues places the *Namao virus* sequence within the clade containing bony fish. Since this virus is known to infect *Acipenser fulvescens* (lake sturgeon) and the particular genome in question was isolated from an infected tissue sample, it is highly likely that this sequence is the result of sample contamination from the host organism. The *Methanophagales archeon* sequence is placed very close to bacterial sequences at a distance which is inconsistent with even the slow divergence of core housekeeping genes shared between bacteria and archaea. As *PADI* does not appear to be part of the core archaeal genome as it is not a housekeeping gene vital for archaeal cell survival, it is unlikely that this sequence would show such a slow rate of evolution while also being lost simultaneously within all other archaeal species.

**Figure S3:**
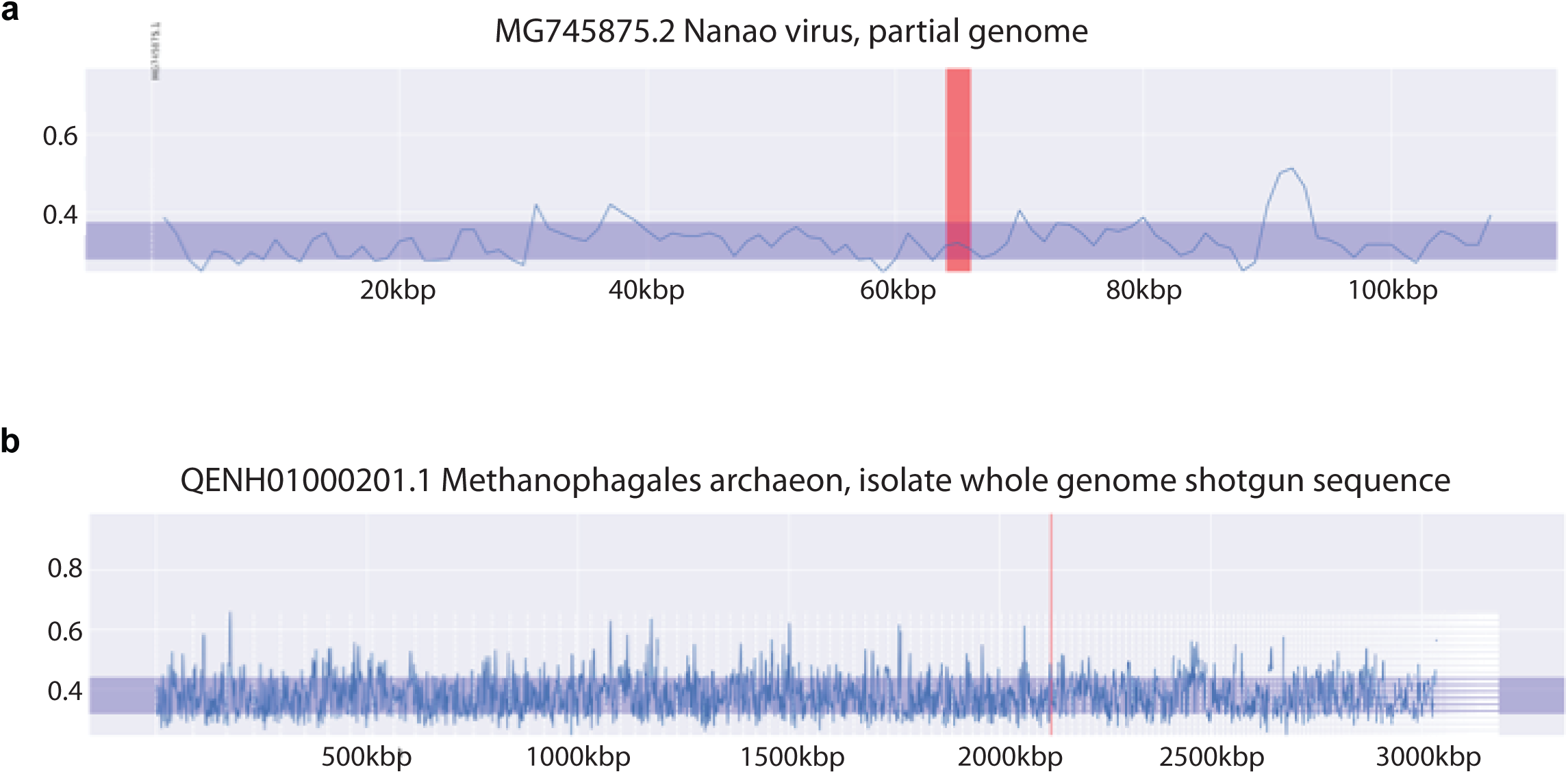
K-mer spectra analysis of genomic contigs of spurious viral (a) and archaeal (b) hits. The norm of the vector defined by the difference between the genomic average and the K-mer spectrum of each window is shown over all contigs in both assemblies. Vertical white dashed lines denote the start of a contig. The horizontal blue shaded region shows the standard deviation around the average distance of all windows relative to the genomic average. The red vertical shaded area shows the position of the putative PADI homologue within the genome. Contig identifiers are not shown for the genome of Methanophagales archeon due to the high number of small contigs. The assembly quality of this genome is fairly poor, contigs are ordered by length and do not represent the actual arrangement. K-mer spectra for contigs shorter than 1kbp were not calculated. Neither of the genomes present a region showing anomalous K-mer spectra typical of a continuous horizontally transferred genomic region. This indicates that the two PADI sequences were falsely attributed to these genomes during assembly due to their K-mer spectra being close to the genomic average of these two organisms by chance.

**Figure S4:**
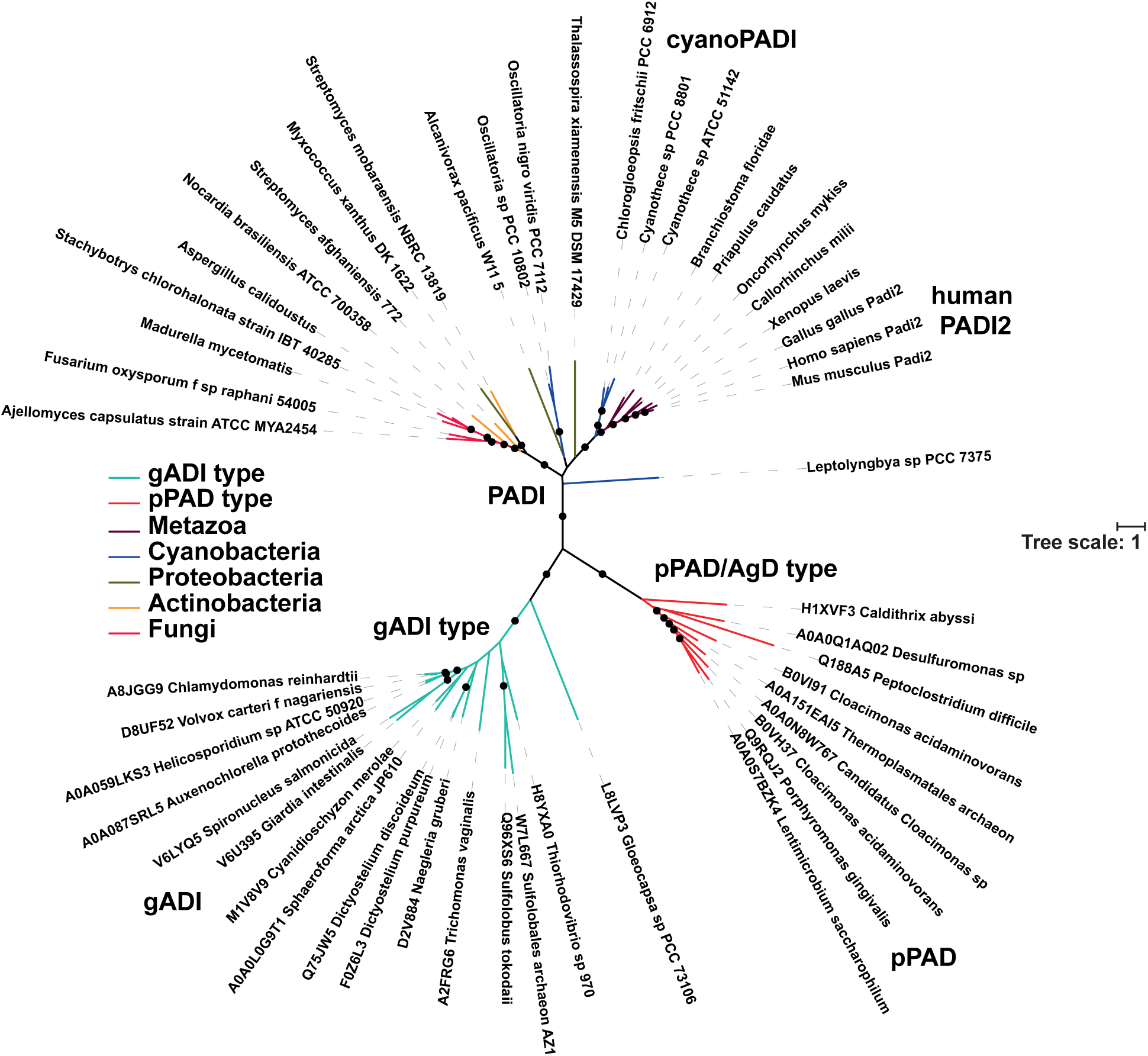
Metazoan PADI sequences are evolutionarily distinct from other bacterial and eukaryotic citrullinating enzymes. PADIs form distinct clades to ADIs, AgDs, gADI (arginine deiminase from Giardia lamblia) and pPAD (porphyromonas-type peptidylarginine deiminase from Porphyromonas gingivalis) sequences. The tree is shown unrooted with solid circles indicating consensus node support of >95%.

**Figure S5:**
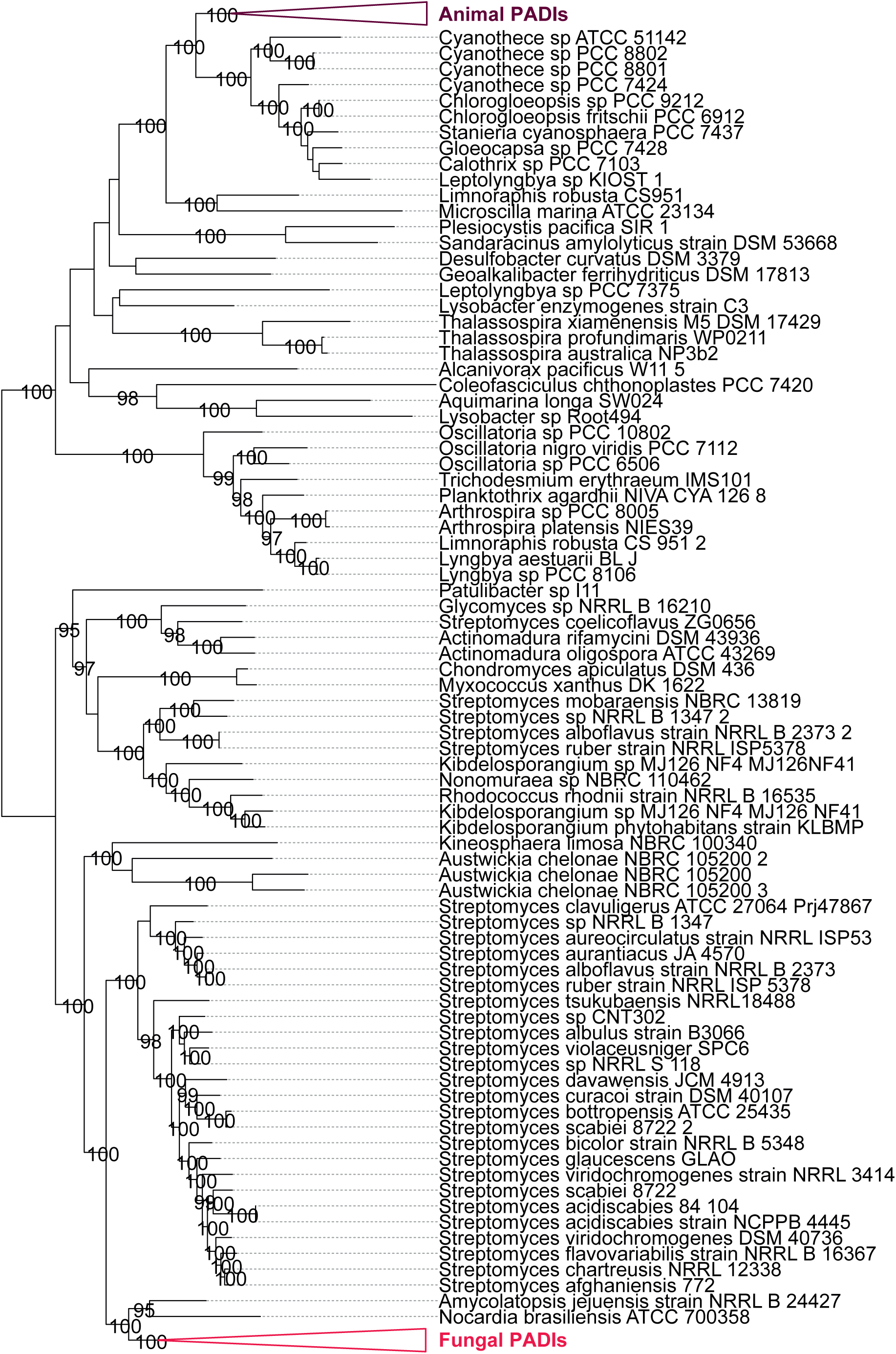
PADI phylogeny. Phylogenetic analysis of all putative bacterial PADI sequences in the PATRIC database and with representative animal and fungal sequences. The tree is shown rooted at the midpoint with solid circles indicating consensus node support of >95% and a number of critical nodes labeled in full.

**Figure S6:**
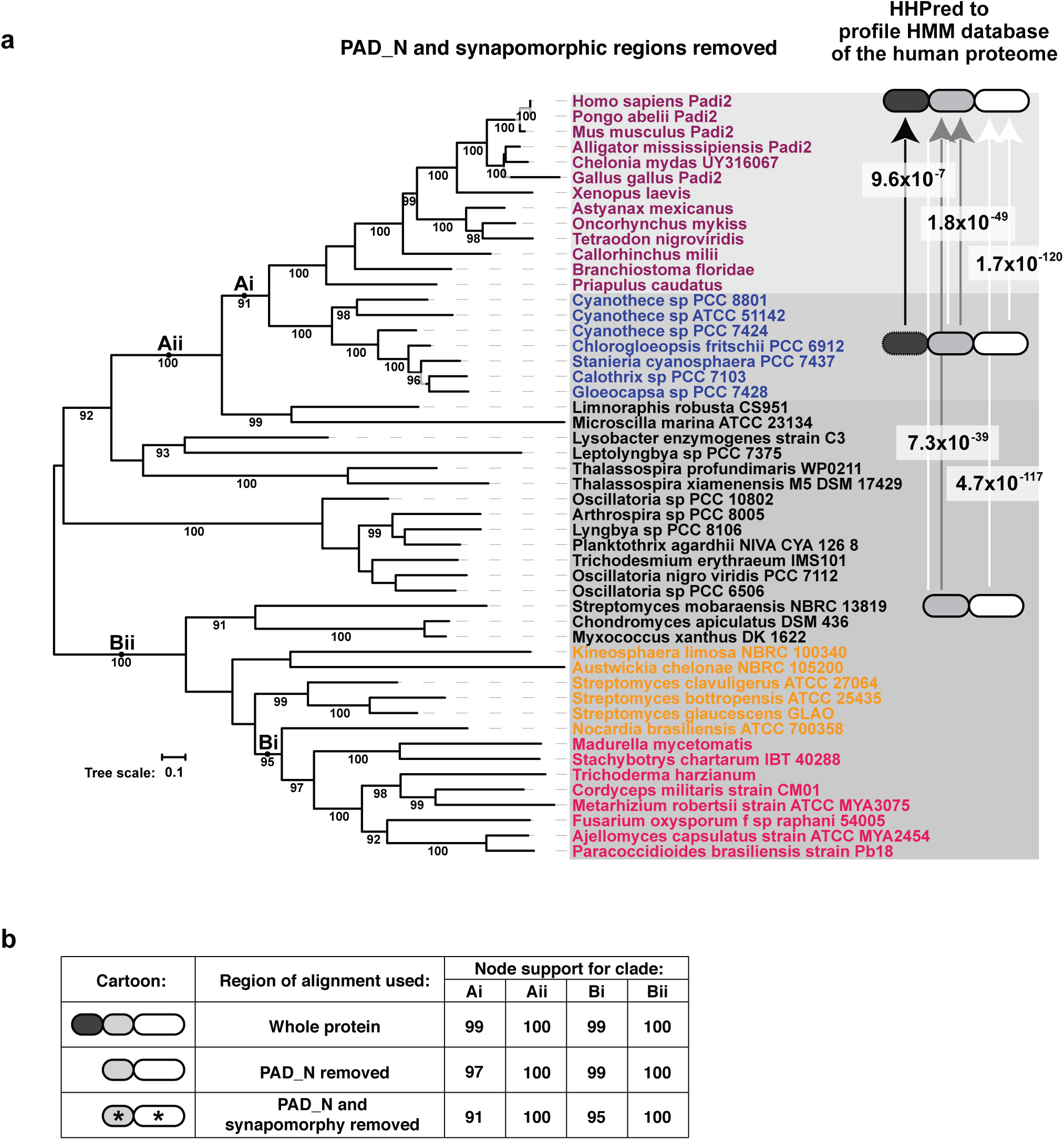
Domain architecture analysis of PADI orthologues. **a)** Representation of PADI domain architecture across the phylogenetic tree. Protein regions aligning to the metazoan PADI domains were extracted from sequences represented in the different clades of the tree (firstly of cyanobacterial sequences and secondly of a mixture of bacterial and fungal sequences). An HMM profile was made and HHPred was used to search against a database of profiles made of the entire human proteome, and against a database of Pfam domain profiles. The E-values are given for these searches where significant sequence similarity could be identified from HHPred searches. **b)** Phylogenetic analyses from Figure 1a (top row) were repeated using an alignment where the PAD_N domain (middle row), or both the PAD_N domain and regions of synapomorphy (bottom row) were removed. Maximum likelihood inference using IQTree was used in all three cases and ModelFinder was used to select the best performing fixed empirical rate matrix (WAG + R5 +F0). Node support values correspond to clades annotated in Figure 1a as topologies were congruent.

**Figure S7:**
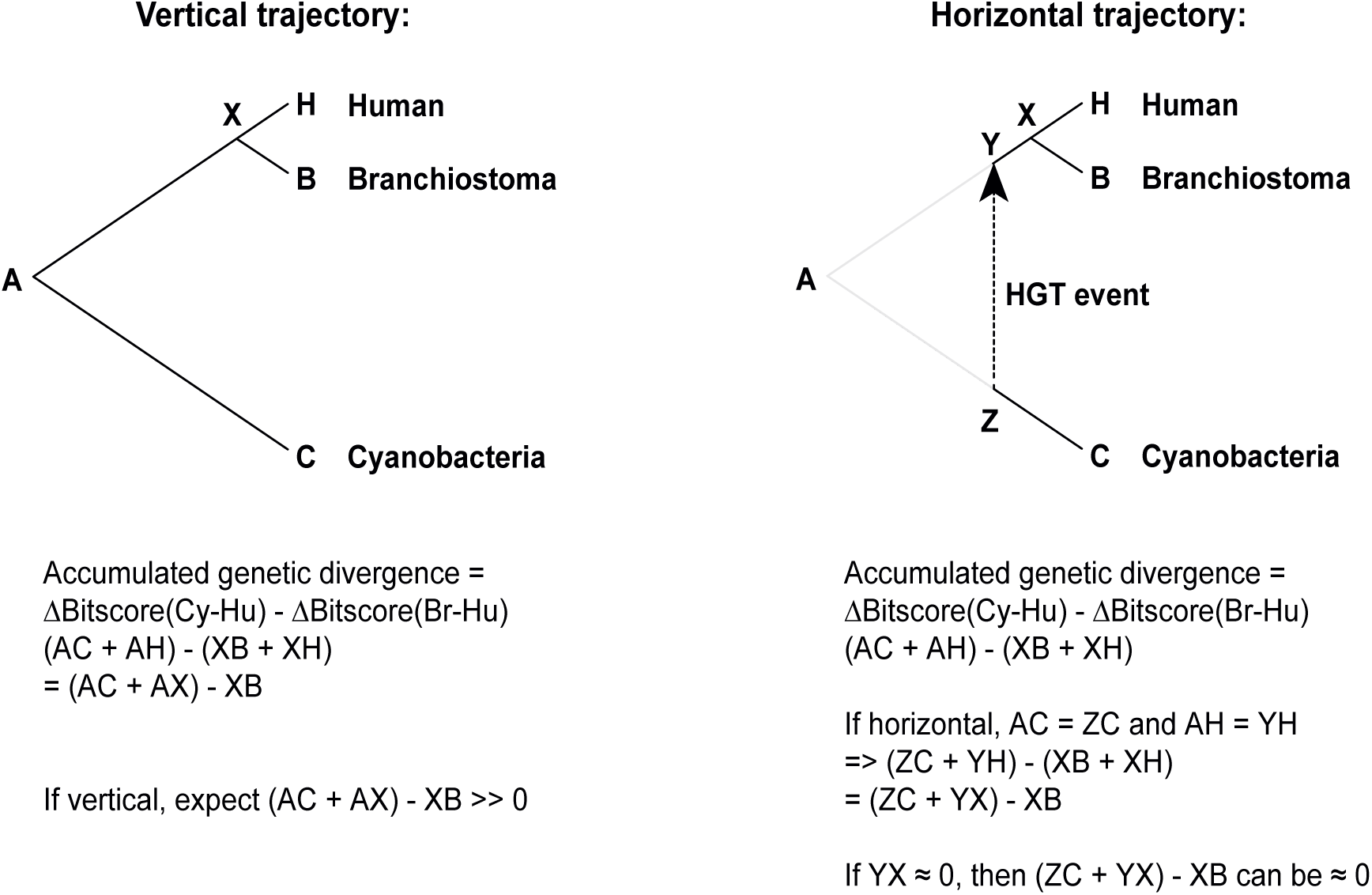
Calculating the AGD of a given protein between its homologues in *Homo sapiens, Branchiostoma spp. and Cyanothece spp*. The bitscore density of the similarity of the cyanobacterial homologue to the human sequence Δbitscore_Cy-Hu_ (AC+AH) and the bitscore density of the similarity of the branchiostomal homologue to the human sequence Δbitscore_Br-Hu_ (XB+XH) are calculated. In a vertical scenario the AGD, given by (AC+AX) – XB, will be much greater than zero. In a horizontal scenario, (ZC+YX) may be approximately equal to XB and so the AGD, given by (ZC + YX) – XB may be close to zero.

**Figure S8:**
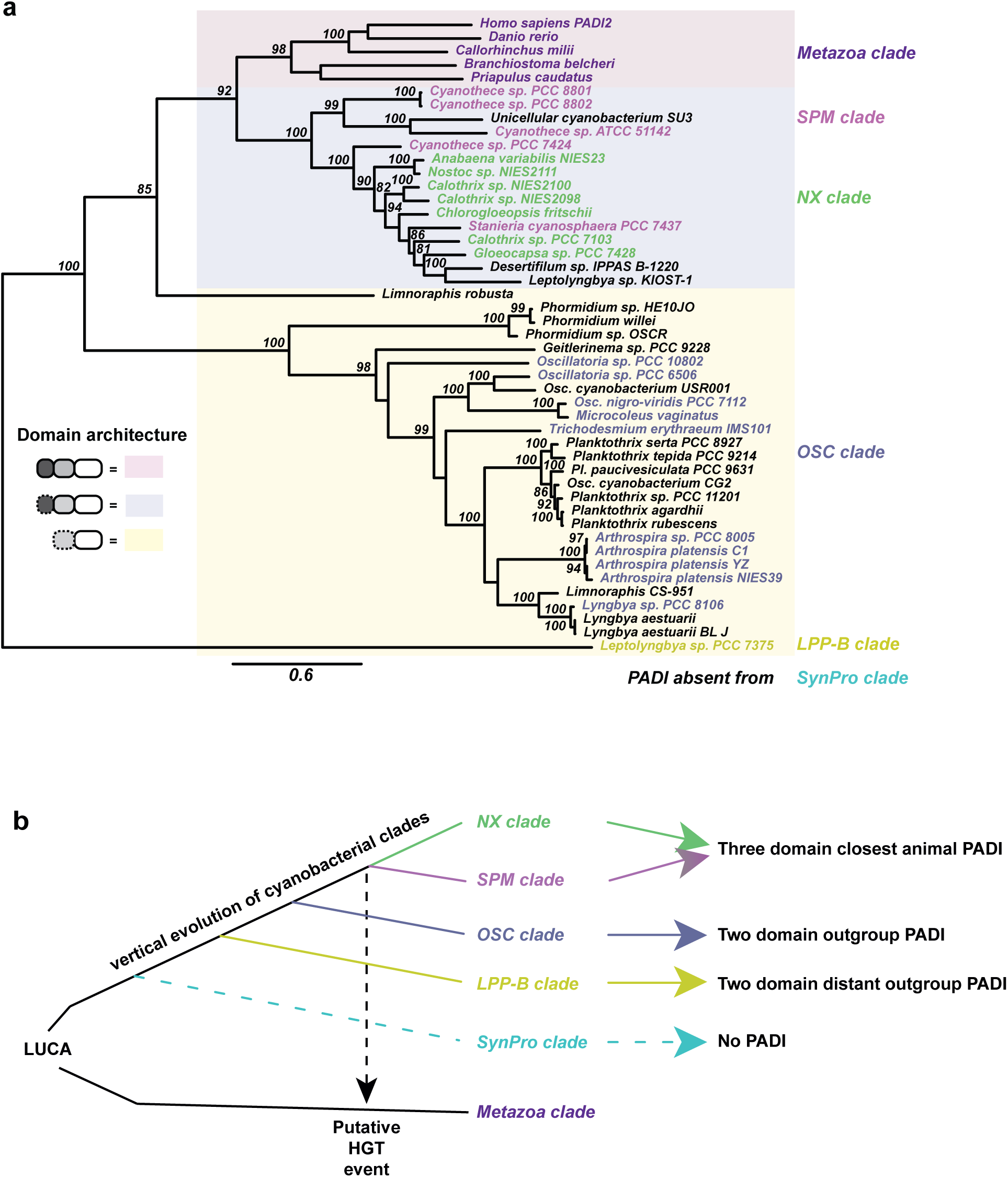
Cyanobacterial origin of the PADI sequence. **a)** Phylogenetic analysis of cyanobacterial PADI sequences reproduces the known clades of cyanobacterial evolution. Colours and names of clades are used as in Uyeda *et al*. Species that were not analysed in Uyeda *et al*. are in black. Sequences were aligned with MUSCLE and the tree was built using PhyML 3.0. Bootstrap support >80% is indicated on branches from 250 bootstrap replicates. The domain architecture of each sequence was analysed and is represented in the legend. **b)** Schematic showing the proposed HGT event from cyanobacteria to metazoa. PADI present in a last common ancestor of the *NX/SPM* clades of cyanobacteria, which possess a three-domain PADI, and was transferred to an ancient last common ancestor in the metazoan lineage.

**Figure S9:**
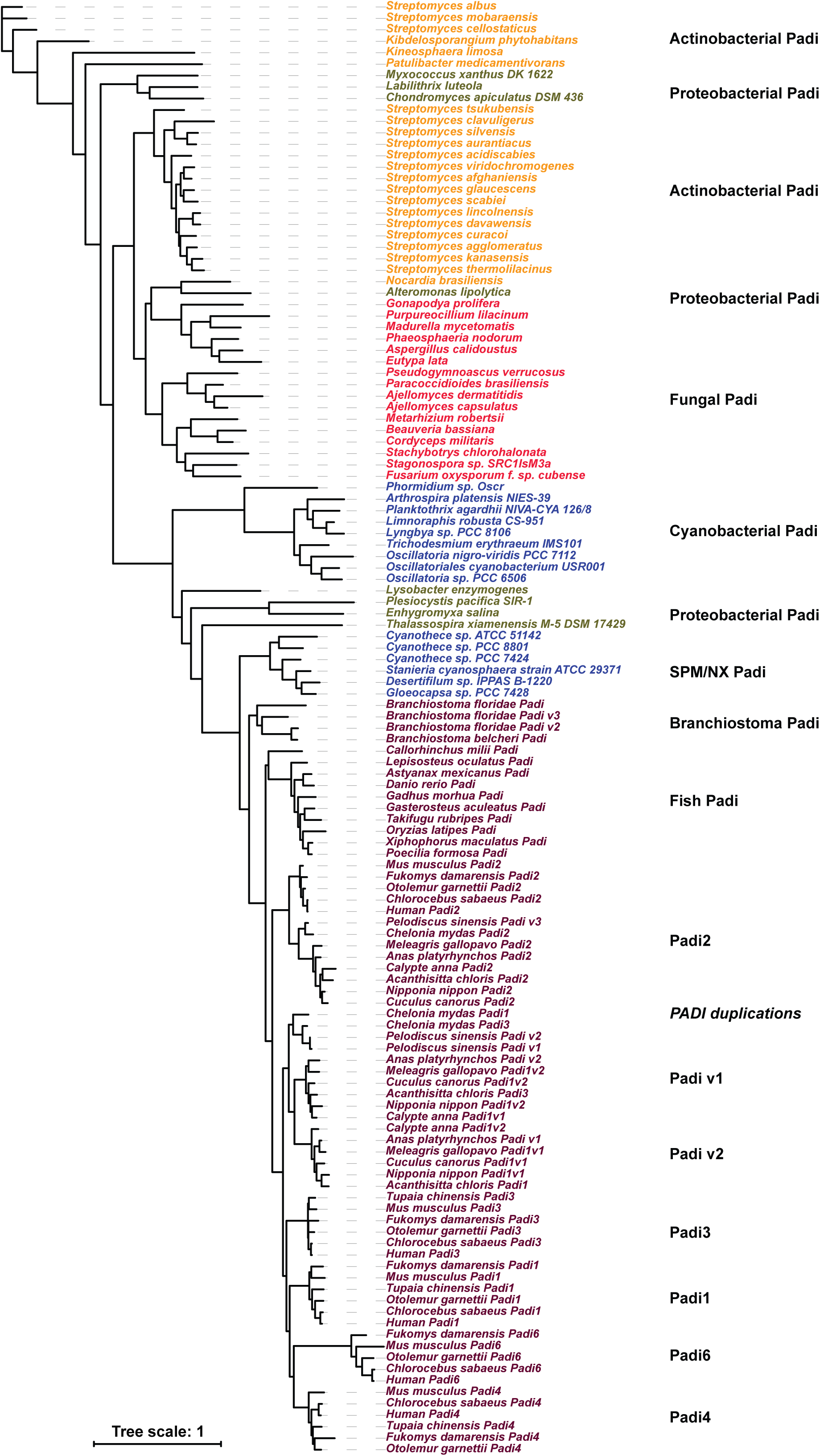
The PADI sequence was retained and underwent multiple duplications in metazoa. Phylogenetic analysis of a large number of putative PADI sequences showing the multiple duplications in metazoa. The first duplication produced two new orthologues present in reptiles and birds which cluster in their own groups denoted PADIv1 and PADIv2. Subsequent duplications in mammals produced the separation of clusters of PADI1, PADI3, PADI4 and PADI6.

